# Extracellular matrix sulfation in the tumor microenvironment stimulates cancer stemness and invasiveness

**DOI:** 10.1101/2023.10.17.562578

**Authors:** Alican Kuşoğlu, Deniz Örnek, Aslı Dansık, Ceren Uzun, Sena Nur Özkan, Sevgi Sarıca, Kardelen Yangın, Şevval Özdinç, Duygu Turan Sorhun, Nuriye Solcan, Efe Can Doğanalp, Øystein Arlov, Katherine Cunningham, İsmail C. Karaoğlu, Seda Kizilel, Ihsan Solaroglu, Pınar Bulutay, Pınar Fırat, Suat Erus, Serhan Tanju, Şükrü Dilege, Gordana Vunjak-Novakovic, Nurcan Tuncbag, Ece Öztürk

## Abstract

Tumor extracellular matrices (ECM) exhibit aberrant changes in composition and mechanics compared to normal tissues. Proteoglycans (PG) are vital regulators of cellular signaling in the ECM with ability to modulate receptor tyrosine kinase (RTK) activation via their sulfated glycosaminoglycan (sGAG) side chains. However, their role on tumor cell behavior is controversial. Here, we demonstrate that PGs are heavily expressed in lung adenocarcinoma patients in correlation with invasive phenotype and poor prognosis. We developed a bioengineered human lung tumor model which recapitulates the increase of sGAGs in tumors in an organotypic matrix with independent control of stiffness, viscoelasticity, ligand density and porosity. Our model reveals that increased sulfation stimulates extensive proliferation, epithelial-mesenchymal transition and stemness in cancer cells. We identified the FAK-PI3K-mTOR signaling axis as a mediator of sulfation-induced molecular changes in cells upon activation of a distinct set of RTKs within tumor-mimetic hydrogels. We demonstrate that the transcriptomic landscape of tumor cells in response to increased sulfation resembles native PG-rich patient tumors through employing integrative omics and network modeling approaches.

## Introduction

The tumor microenvironment (TME) is a dynamic niche in which tumor cells and a plethora of stromal and immune cell types interact within tumor-specific extracellular matrix (ECM), and plays a fundamental role in regulating signaling events involved in tumorigenesis and dissemination^1^. The biochemical and mechanical characteristics of ECM are deregulated during malignancy, resulting in activation of various cellular mechanisms that partake in tumor growth, angiogenesis, metastasis, and immune suppression^2^. Dissecting the effect of aberrant changes in the ECM on tumor cell behavior is crucial for a deeper understanding of the complex orchestration of signaling events that govern disease progression. This calls for human tumor models with ability to recapitulate the key aspects of the ECM and enable their controlled tunability. The need for such models has led to an intersection of tissue engineering and cancer research. Biological materials such as reconstituted basement membrane (rBM)^3^ and collagen^4^ as well as synthetic polymers such as polyethylene glycol (PEG)^5–8^ have become gold standard tools for three-dimensional (3D) *in vitro* modeling of tumor tissues which established the importance of replicating tumor mechanics and composition.

Tumors are marked by distinct expression of cell instructive ECM ligands compared to native organs where they emerge^9^. Proteoglycans (PGs) are heterogenous glycoproteins with sulfated glycosaminoglycan (sGAG) side chains that have vital functions in the ECM^10^. Due to their negatively charged sulfate moieties, sGAGs infer affinity to bioactive ligands, control their availability and mediate formation of ternary complexes with ligands and receptor tyrosine kinases (RTKs) which lead to their activation and downstream signaling cascades^10^. RTK family receptor signaling has key roles in many stages of tumor progression and most cancers are characterized with driver mutations in RTKs^11^. Expectedly, many types of tumor tissues demonstrate increase in sGAG content and altered sulfation pattern^12^. Although deregulation of sGAG biosynthesis and post-translational modification have been shown to correlate with poor prognosis, the effect of increase in sGAGs on tumor growth and invasion has been controversial^12–15^. Studies have reported both tumor promoting and inhibiting effects of sGAG supplementation^13–18^. The use of sGAGs has even been proposed as a therapeutic approach in cancer^19^. These studies strongly suggest the need for an unprecedented 3D human tumor model with ability to present sGAGs to tumor cells within their microenvironmental milieu and allow recapitulation of their aberrant increase while having control over ECM content, stiffness, viscoelasticity and network porosity to elucidate their distinct effects on tumor cells.

To investigate the effect of increased sulfation within an engineered model, a representation of the heathy tissue ECM onto which malignant characteristics could be controllably introduced is required. However, materials such as rBM, which is derived from Engelbreth-Holm-Swarm (EHS) sarcoma and has an undefined composition with high variability, lack a faithful recapitulation of healthy ECM^3^. Decellularization of native organs offer the advantage of preserving the tissue-specific composition of ECM^20^. We have recently developed bovine-derived decellularized lung ECM hydrogels with low batch-to-batch variability and high compositional resemblance to human lungs^21^. On the other hand, the use of native sGAGs as well as sGAG-mimetic materials have been pursued in tissue engineering^5,22–23^. Alginate, an inert biopolymer, can be sulfated to successfully mimic native sGAGs in both exerting affinity to growth factors and mediating their interactions with RTKs^22–23^. Precise tunability of sulfation and enabling of the alginate backbone for mechanical modulation renders this approach advantageous over the use of native sGAGs.

Here, we introduce a biomaterial-based 3D human lung tumor model constituted of double-network hydrogels of decellularized lung ECM with sGAG-mimetic alginate sulfate to investigate the effect of sulfation in the TME on growth and progression of lung tumor cells. Addressing the aforementioned challenges, our model allows tunability of sulfation within an organotypic network while enabling control over ligand density and tissue mechanics. Our data demonstrates that increased sulfation acts as an essential regulator in the lung TME which promotes growth, stemness and epithelial-mesenchymal transition (EMT) via the RTK-phosphatidylinositol 3-kinase (PI3K) signaling axis.

## Results

### Elevated proteoglycan expression correlates with invasiveness and poor survival in lung adenocarcinoma patients

Non-small cell lung adenocarcinoma (LUAD) is the most common type of lung cancer, the leading cause of cancer-related mortality worldwide^24^. To validate our hypothesis of sulfation-mediated regulation of tumor progression, we initially performed bioinformatics analyses on a 510 patient LUAD cohort to reveal differential expression of PGs in patient tumors. Analyses of RNA sequencing (RNAseq) data derived from The Cancer Genome Atlas (TCGA) for 34 PG-encoding genes in LUAD samples relative to normal lung tissue demonstrated that the expression of the majority of PG genes was increased in patient tumors (Fig. 1a). We then defined two patient groups within the cohort. PG positive (PG+) group included patients which had z-score equal to or higher than a cut-off value of three for at least three PG genes, whereas PG negative (PG-) group had zero genes. PG+ group represented 48% of all patients, had more than five-fold higher number of patients, significantly higher PG score and a poorer survival curve compared to the PG-group (Fig. 1b-d). EMT in lung cancer cells highly correlates with poor prognosis^25^. We next calculated the EMT score for all patients from the RNAseq data (see Methods). The correlation of EMT score and PG score in LUAD patients was strikingly high (ρ = 0.771) (Fig. 1e). EMT and PG scores of patients were also calculated for protein expression from CPTAC data which similarly showed a significantly high correlation (ρ = 0.457) (Fig. 1f). We then collected surgically resected tumor and matched healthy parenchyma tissues from 5 LUAD patients to perform sGAG quantification and histopathological assessments. Patient-derived tumor tissues had significantly higher amounts of sGAGs (2.85-fold) than normal tissues (Fig. 1g). Consistently, Alcian blue staining revealed increased sGAG deposition in tumor tissues (Fig. 1h). Together, these results confirmed the relevance of biomimicking the increased sulfation of lung tumor matrices with an engineered human tumor model.

**Figure 1.**
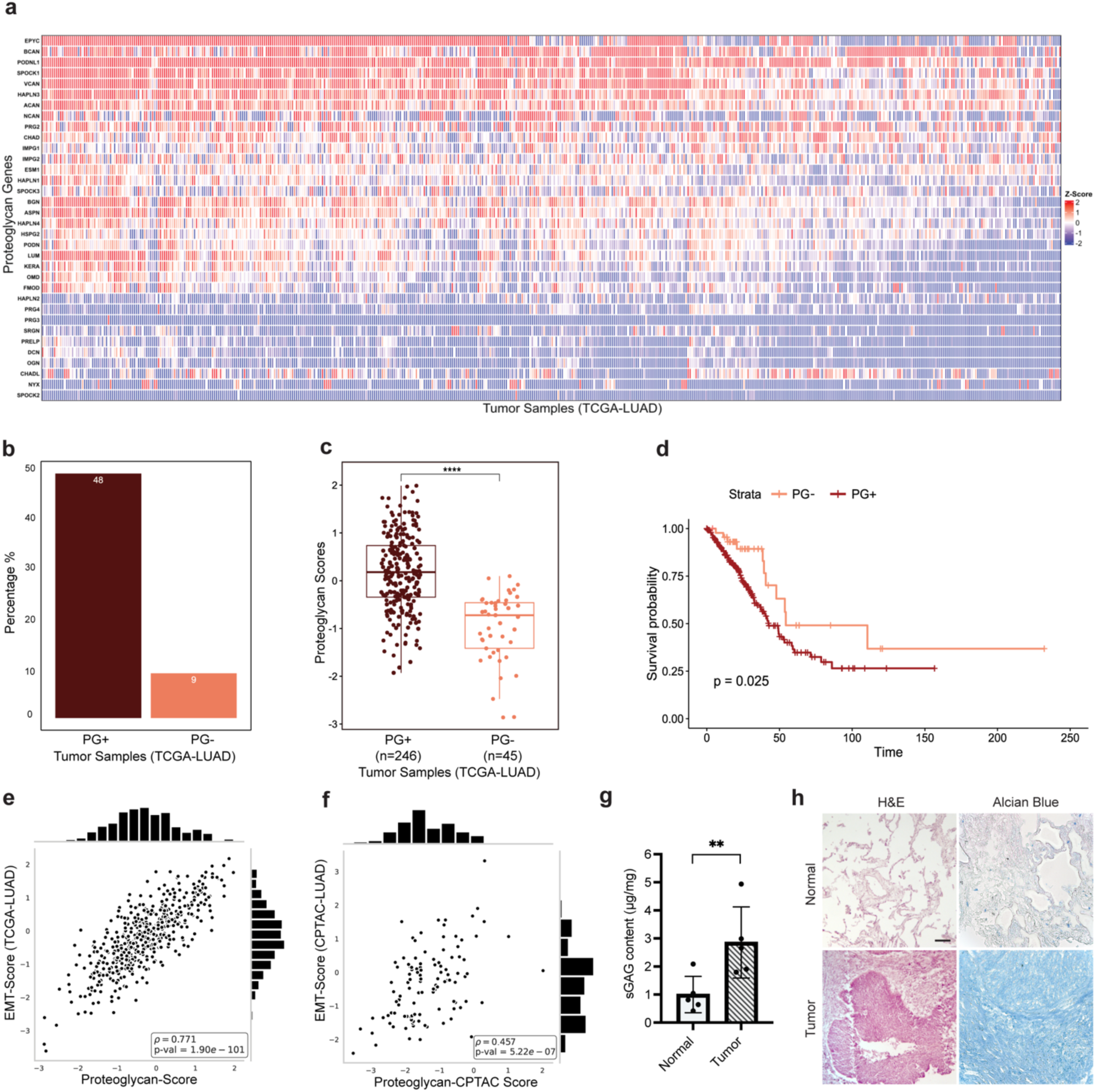
Elevated proteoglycan expression correlates with invasiveness and poor survival in LUAD patients. **a** Hierarchical clustering heatmap of proteoglycan (PG) mRNA expression z-scores in LUAD patient tumors relative to normal tissue samples where the rows represent PG genes and columns indicate patients. **b** Percentages of PG+ and PG-tumor samples in TCGA LUAD patient cohort. A sample is considered PG+ if z-score is greater than the cut-off of three for at least three genes, while a sample is considered PG-if the z-score is higher than the cut-off, but the number of genes is equal to zero. **c** Box plot comparison of PG scores of PG+ and PG-patient groups. Each box extends from lower to the upper quartile where the horizontal line indicates the median value. Each dot represents the PG score of an individual patient. Statistical analysis was performed with Wilcoxon test, p = 1.5e^-14^. **d** Survival plots of PG+ and PG-LUAD patients. Log-rank test was used to assess significance, p = 0.025. **e** Regression analysis on epithelial-mesenchymal transition (EMT) scores and proteoglycan scores using the mRNA (p = 1.9e^-101^) and **f** protein expressions (p = 5.22e^-07^). Spearman rank correlation was used to calculate the correlation values (ρ). **g** Sulfated glycosaminoglycan (sGAG) quantification in LUAD patient-derived tumor and matched normal lung samples, n = 5 biological replicates. Data is represented as mean ± S.D and statistical significance was analyzed using a paired, two-tailed student’s t-test, p < 0.01. **h** Hematoxylin and eosin (H&E) and Alcian blue staining of normal lung parenchyma and tumor tissue (scale bar: 100 μm).

### Mimicking the increased sulfation in the TME drives aberrant proliferation of lung tumor cells

To model the increase of sulfation in the TME, decellularized and reconstituted lung-derived ECM (dLung), representing the matrix composition at the tumor’s site of origin, was combined with either alginate (Alg) or sGAG-mimetic alginate sulfate (S-Alg) to obtain interpenetrating, double-network hydrogels (Fig. 2a). Decellularization of bovine lungs was carried out following our freeze-thaw method^21^ and validated for elimination of nuclear content, preservation of ECM constituents, gelation and biocompatibility (Fig. 2a, Supplementary Fig. 1). Degree of sulfation (DS) of alginate can be tuned without altering structural features of the polymer such as monosaccharide structure, chain confirmation and flexibility^22–23^. Sulfation of alginate was performed to yield a DS value of 0.41 sulfate groups per monomer (Supplementary Fig. 2). Tumor-mimetic hydrogels were fabricated with dLung and S-Alg, S-AlgLung, whereas healthy-mimetic hydrogels, AlgLung, comprised of unmodified alginate. Both hydrogels had equal concentration of dLung (5,4 mg/ml) and alginates (10 mg/ml). Alginate and derivatives, due to their ionic crosslinking capability in the presence of divalent cations allow tunability of stiffness independently from ECM composition and porosity^26^. We modulated crosslinker (Ca^2+^) density to match the stiffness of the tumor-mimetic (S-AlgLung) and healthy-mimetic (AlgLung) hydrogels which demonstrated similar storage moduli and loss tangent with oscillatory rheology (Fig. 2b, c). The stiffness range for the hydrogels was tuned to match the stiffness of healthy lung tissue, whose Young’s modulus was reported in 1-2 kPa range^27^. This was aimed to investigate the sole effect of increased sulfation in the cellular microenvironment without the effect of stiffening. PGs have been shown to interact with mechanosensitive receptors such as integrin family to induce distinct synergistic signaling pathways^28^. Apart from stiffness, viscoelasticity has been shown to be an important parameter which affects cancer cell behavior. Modulation of molecular weight has been reported as a means to tune plasticity of alginate hydrogels^29^. Sulfation caused a significant decrease in the molecular weight of alginate as indicated by size exclusion chromatography without compromising its hydrogel forming capacity (Fig. 2d, Supplementary Table 1). Therefore, we investigated the stress-relaxation behavior of AlgLung and S-AlgLung hydrogels with a creep-recovery test. Surprisingly, the two hydrogels demonstrated very similar relaxation curves and permanent strain values (Fig. 2e, f). Next, we assessed porosity in the hydrogels with dextran release assay using two different sizes (10 kDa and 70 kDa) which showed similar release curves for both AlgLung and S-AlgLung (Fig. g, h). Thus, our model allowed alteration of sulfation in the presence of organotypic ECM independently from ligand density, hydrogel stiffness, plasticity and porosity.

To investigate the effects of increased sulfation on cellular proliferation, we encapsulated LUAD-derived A549 cells into AlgLung and S-AlgLung hydrogels and cultured up to 4 weeks. S-AlgLung hydrogels stimulated extensive growth of A549 cells starting from the first week (Fig. 2i, j). Cell viability was not compromised with unmodified alginate in AlgLung hydrogels but rather cell growth was limited (Supplementary Fig. 3). Moreover, when cells were cultured in double-network hydrogels of alginate and rBM (Matrigel), AlgMat, potent cell growth was again observed demonstrating the growth-limiting effect of organotypic lung matrix (Supplementary Fig. 4). AlgMat hydrogels had over 2-fold higher amount of sGAGs than AlgLung (Supplementary Fig. 5), however, the undefined, tumor-derived composition of Matrigel does not allow the attribution of observed phenomena to a single component. This further emphasizes the advantage of using healthy tissue-derived matrices and introduction of malignant ECM parameters in a controllable fashion to model their effects on cancer cells. Cellular cluster formation in hydrogels was analyzed which exhibited a significant increase in cluster number and area in S-AlgLung hydrogels as well as a decrease in cluster circularity indicating a more invasive phenotype (Fig. 2k, l, m). In line with invasiveness, collective cell migration was observed at the periphery of S-AlgLung hydrogels at week 4, suggesting that the sulfated microenvironment both promoted cluster formation and facilitated migration (Supplementary Fig. 6). We next stained cells with histopathological markers^30^ for non-small cell lung cancer (NSCLC), thyroid transcription factor (TTF-1) and tumor protein p63 (Fig. 2n). Cells in S-AlgLung hydrogels stained positive for TTF-1, a common lineage marker for diagnosis, which is significantly increased in lung tumors compared to normal tissue^31^. p63, a characteristic marker for squamous carcinoma, is found in a subset of adenocarcinomas and is a potent regulator of proliferation and cluster formation^32^. Recently, TTF-1/p63 double-positive LUAD tumors have been dubbed as a basal-like and aggressive phenotype^33^ in line with the proliferative and invasive phenotype we observe in S-AlgLung hydrogels.

**Figure 2.**
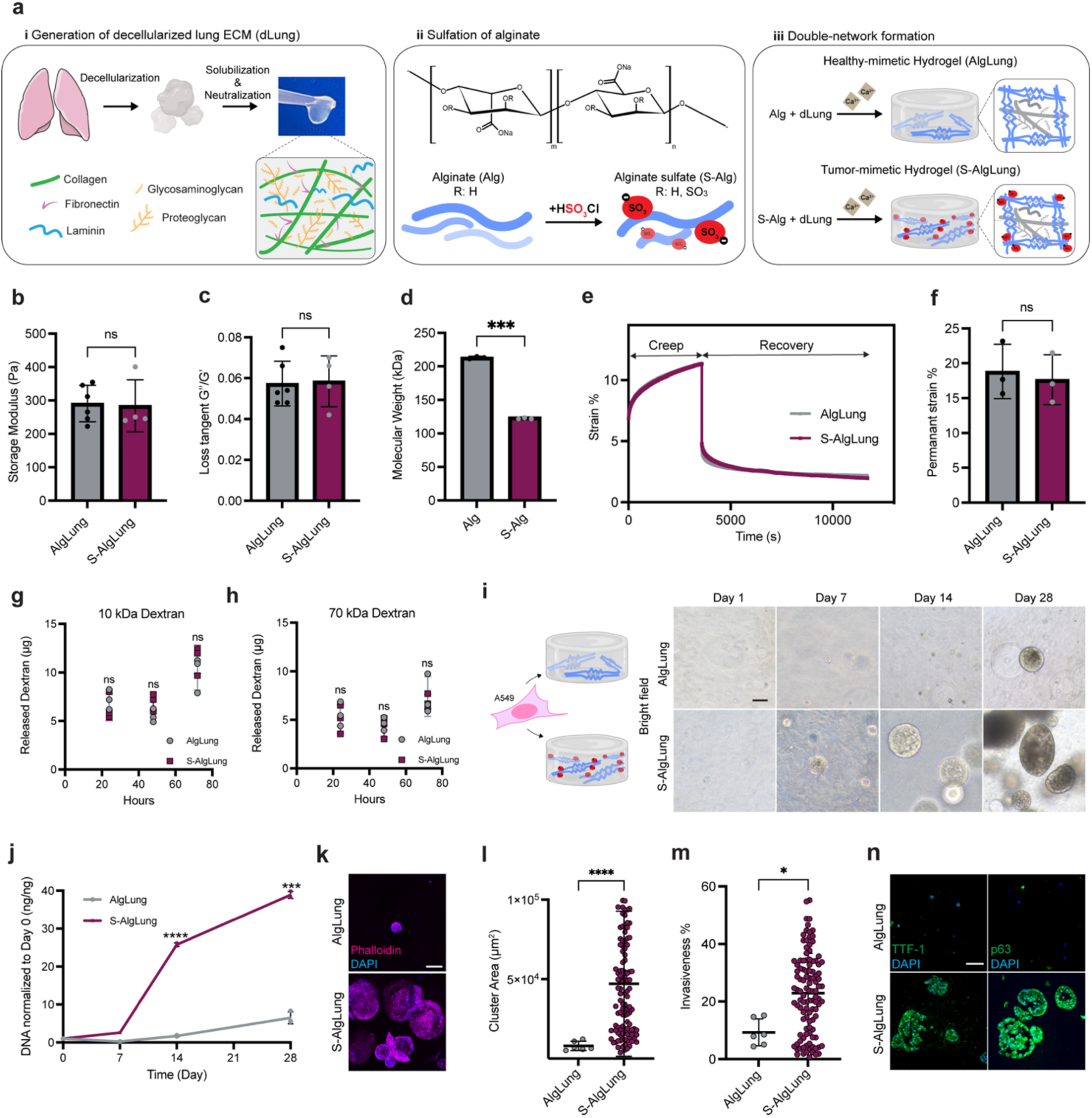
Mimicking the increased sulfation in the TME drives aberrant proliferation of lung tumor cells. **a** Schematic illustration of the fabrication of AlgLung and S-AlgLung hydrogels using decellularized lung ECM and alginate/alginate sulfate. **b** Storage modulus and **c** Loss tangent (G’’/G’) of AlgLung and S-AlgLung hydrogels, ns not significant. **d** Molecular weight of alginate (Alg) and alginate sulfate (S-Alg) characterized by SEC-MALS, p < 0.001. **e** Creep and Recovery test of AlgLung and S-AlgLung hydrogels. **f** Permanent strain of AlgLung and S-AlgLung hydrogels obtained from creep and recovery tests, ns not significant. **g** 10 kDa and **h** 70 kDa dextran release from AlgLung and S-AlgLung hydrogels over 72 hours, ns not significant. **i** Bright field images of A549 cells in AlgLung and S-AlgLung hydrogels at day 1, 7, 14, and 28. **j** Quantification of DNA content in AlgLung and S-AlgLung hydrogels normalized to day 0, **p < 0.01, ***p < 0.001. **k** Immunofluorescence image of A549 clusters stained for Phalloidin (magenta) and DAPI (blue) in AlgLung and S-AlgLung hydrogels (scale bar: 100 μm). **l** Quantification of cluster area (μm^2^) and **m** invasiveness (%) of cells grown in AlgLung and S-AlgLung hydrogels at day 28, *p < 0.05, ****p < 0.0001. **n** Immunofluorescence staining of lung adenocarcinoma markers TTF-1 and p63 (green) and DAPI (blue) in A549 cells grown in AlgLung and S-AlgLung hydrogels (scale bar: 100 μm). **b-m** Quantitative data is represented as mean ± S.D and statistical significance was analyzed using an unpaired, two-tailed student’s t-test.

### Increased sulfation in the ECM modulates EMT and cancer stemness

sGAGs regulate RTK signaling which in turn activates signaling routes that promote EMT, tumor cell motility and metastasis^11–14^. Therefore, we explored the EMT process that might induce an invasive phenotype in sulfated hydrogels. Immunofluorescence (IF) staining revealed elevated expression of mesenchymal markers including N-cadherin, vimentin, and fibronectin in S-AlgLung hydrogels (Fig. 3a). Consistently, expression of CDH2 (N-cadherin), vimentin and FN1 (fibronectin) genes were significantly increased in sulfated hydrogels (Fig. 3b). In contrast, we also observed an upregulation of epithelial marker E-cadherin on both protein (Fig. 3a) and gene (CDH1) (Fig. 3b) levels in sulfated hydrogels, indicating a lack of conventional “cadherin-switch”^34^ but a more dynamic transition between epithelial-mesenchymal states normally observed in metastatic tumor cells^35^. Such complex, co-expression of these markers is not atypical for clinical LUAD samples^36^ which was recapitulated in tumor-mimetic hydrogels. Besides, E-cadherin is a promoter of spheroid formation correlating with the increased number of clusters in sulfated hydrogels which exhibit localization of E-cadherin in the periphery (Fig. 3a). Moreover, sulfated hydrogels strongly induced the expression of several EMT-regulating transcription factors such as ZEB1, ZEB2, and SNAIL (Fig. 3b), further emphasizing the role of ECM sulfation (Fig. 3c).

Recent studies suggest that secretory mucins MUC5AC and MUC5B are aberrantly expressed in tumor tissues and associated with distant metastases and poor survival in LUAD patients^37^. MUC5B staining reveals enhanced deposition in S-AlgLung hydrogels (Fig. 3d) in line with significantly increased expression of MUC5AC and MUC5B genes (Fig. 3e). Interestingly, studies showed that activation of EMT process can increase cancer stem cell (CSC) population with self-renewal capacity within tumors^38^. Furthermore, a correlation between CSC phenotype and expression of several mucins have been reported^39^. Thus, we next investigated whether sulfated ECM could stimulate stemness in cancer cells. Expression of SOX2, an important stemness marker and regulator of EMT program, was potently increased in S-AlgLung hydrogels (Fig. 3f). Similarly, gene expression of stemness markers SOX2, KLF4, OCT3/4, CD44 and CD133 exhibited strong upregulation upon sulfation (Fig. 3g). sGAGs have been proposed to have a role in regulation of stemness through modulation of WNT/beta-catenin pathway^40^. Consistently, beta-catenin also demonstrated higher expression in sulfated hydrogels (Fig. 3f). We then investigated the correlation of PG expression and stemness (CSC-score) in the TCGA LUAD cohort which revealed a significant correlation (ρ = 0.457) (Fig. 3h). Overall, these findings further validate that increased sulfation in lung tumors promotes an aggressive, mucinous and stem-like phenotype.

**Figure 3.**
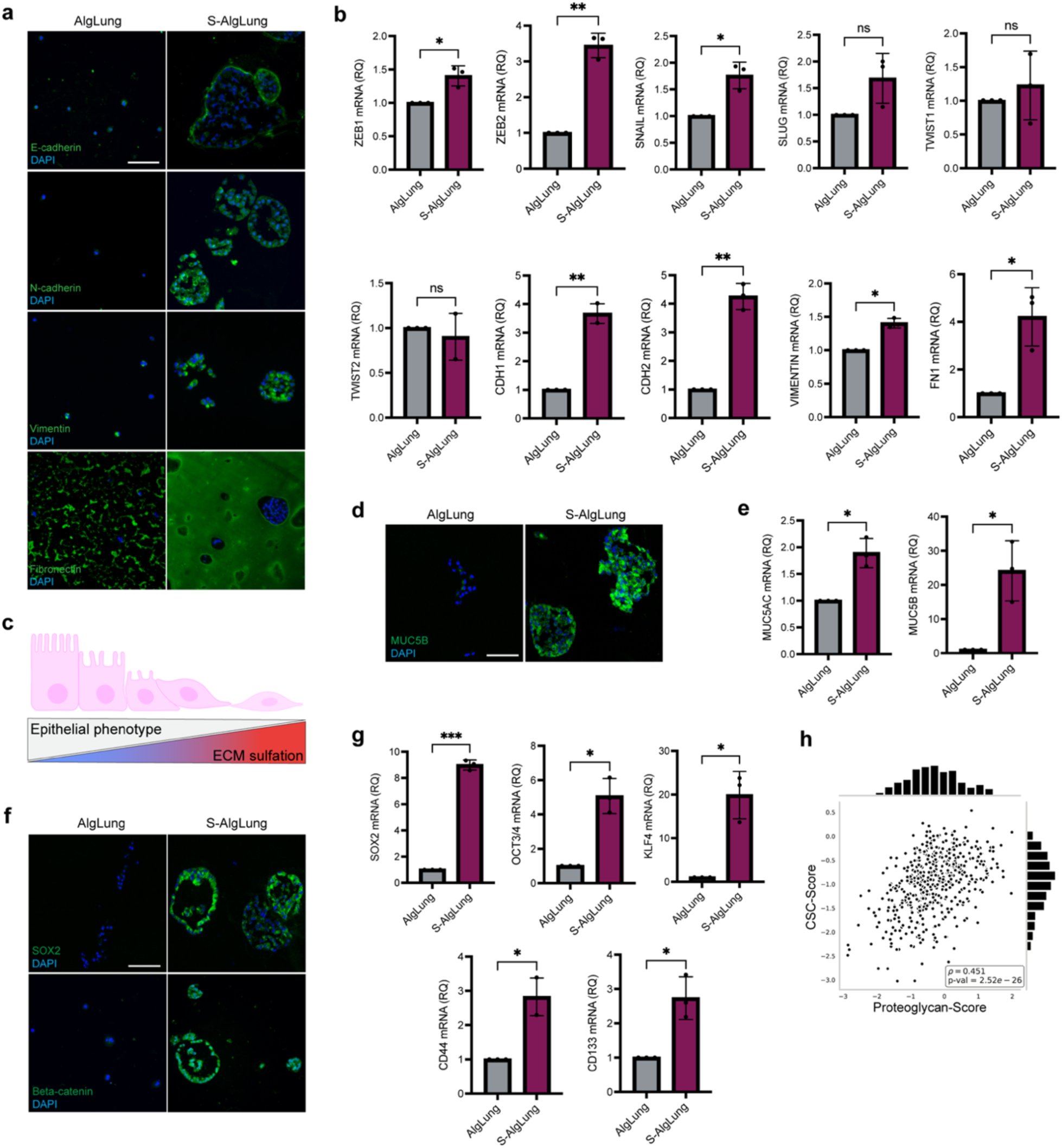
Increased sulfation in the ECM modulates EMT and cancer stemness. **a** Immunofluorescence staining for E-cadherin, N-cadherin, vimentin and fibronectin (green, top to bottom) and DAPI (blue) in A549 cells grown AlgLung and S-AlgLung hydrogels (scale bar = 100 μm) **b** mRNA expression of EMT regulators and markers in A549 cells grown in AlgLung and S-AlgLung hydrogels, ns not significant, *p < 0.05, *p < 0.01. **c** Schematic illustration showing the inverse correlation between epithelial phenotype and extracellular matrix (ECM) sulfation **d** Immunofluorescence staining of MUC5B (green) and DAPI (blue) in A549 cells grown in AlgLung and S-AlgLung hydrogels (scale bar: 100 μm) **e** mRNA expression of MUC5AC and MUC5B in A549 cells grown in AlgLung and S-AlgLung hydrogels, *p < 0.05 **f** Immunofluorescence staining of SOX2 (top) and beta-catenin (bottom) (green) and DAPI (blue) in A549 cells grown in AlgLung and S-AlgLung hydrogels (scale bar: 100 μm) **g** mRNA expression of stemness markers in A549 cells grown in AlgLung and S-AlgLung hydrogels, *p < 0.05, ***p < 0.001 **h** Regression analysis of cancer stem cell (CSC) genes and proteoglycans using the mRNA expression scores in TCGA LUAD patient cohort. Spearman rank correlation was used to calculate the correlation values (ρ) (p = 2.52e^-26^). **b, e, g** Relative quantification (RQ) was used with normalization to AlgLung samples. Data is represented as mean ± S.D and statistical significance was analyzed using an unpaired, two-tailed student’s t-test.

### Receptor tyrosine kinase signaling mediates sulfation-induced tumorigenic phenotype

Next, we wanted to uncover the molecular pathways mediating the response of lung tumor cells to sulfated ECM. sGAGs modulate the common RTK pathways through both increased retention of ligands and enabling ligand-receptor complexes that leads to receptor activation^10^. Epidermal growth factor receptor (EGFR) is overexpressed in more than 25% of lung adenocarcinoma cases which correlates with poor prognosis. Integrin β1 was suggested to be required for the ligand-medicated activation of EGFR^41^. We thus examined EGFR and integrin β1 expression of lung tumor cells in AlgLung and S-AlgLung hydrogels. Both receptors were positively stained on the cell membrane of A549 cells in S-AlgLung hydrogels, suggesting their activation as a possible mediator of sulfation-mediated events (Fig. 4a). However, inhibition of neither receptor was able to block sulfation-induced tumor cell growth in sulfated gels (Fig. 4b, c). We then blocked other RTKs including FGFR and TGFβR. Only blocking FGFR resulted in inhibition of cell growth in S-AlgLung hydrogels and further blocking EGFR and TGFβR did not further enhance the inhibitory effect (Fig. 4d,e) (Supplementary Fig. 7). We performed a phospho-RTK array (49 human RTKs) to validate the activation of FGFR and screened for other possible RTKs which might be activated upon interaction with sulfated ECM (Fig. 4f, g). Phosphorylation of FGFR3, an RTK recently reported to be highly expressed in NSCLC samples^42^ but not FGFR1 was induced in S-AlgLung hydrogels. Interestingly, phosphorylated EGFR was even reduced in sulfated gels (Supplementary Fig. 8). Activation of RYK, a receptor with regulatory role in Wnt/beta-catenin signaling^43^, was also induced in S-AlgLung in line with expression of beta-catenin (Fig. 3f, 4g). After confirming the activated RTKs, we next sought to elucidate the downstream signaling route that controls the sulfation-induced effects. We treated A549 cells in S-AlgLung hydrogels with small molecule inhibitors targeting the PI3K-Akt-mTOR, CDC42-NWASP-Arp2/3 as well as FAK signaling axes. Inhibition of PI3K, mTOR as well as FAK completely abrogated the sulfation effect on cell growth without compromising viability (Fig. 4h-j) (Supplementary Fig. 9).

**Figure 4.**
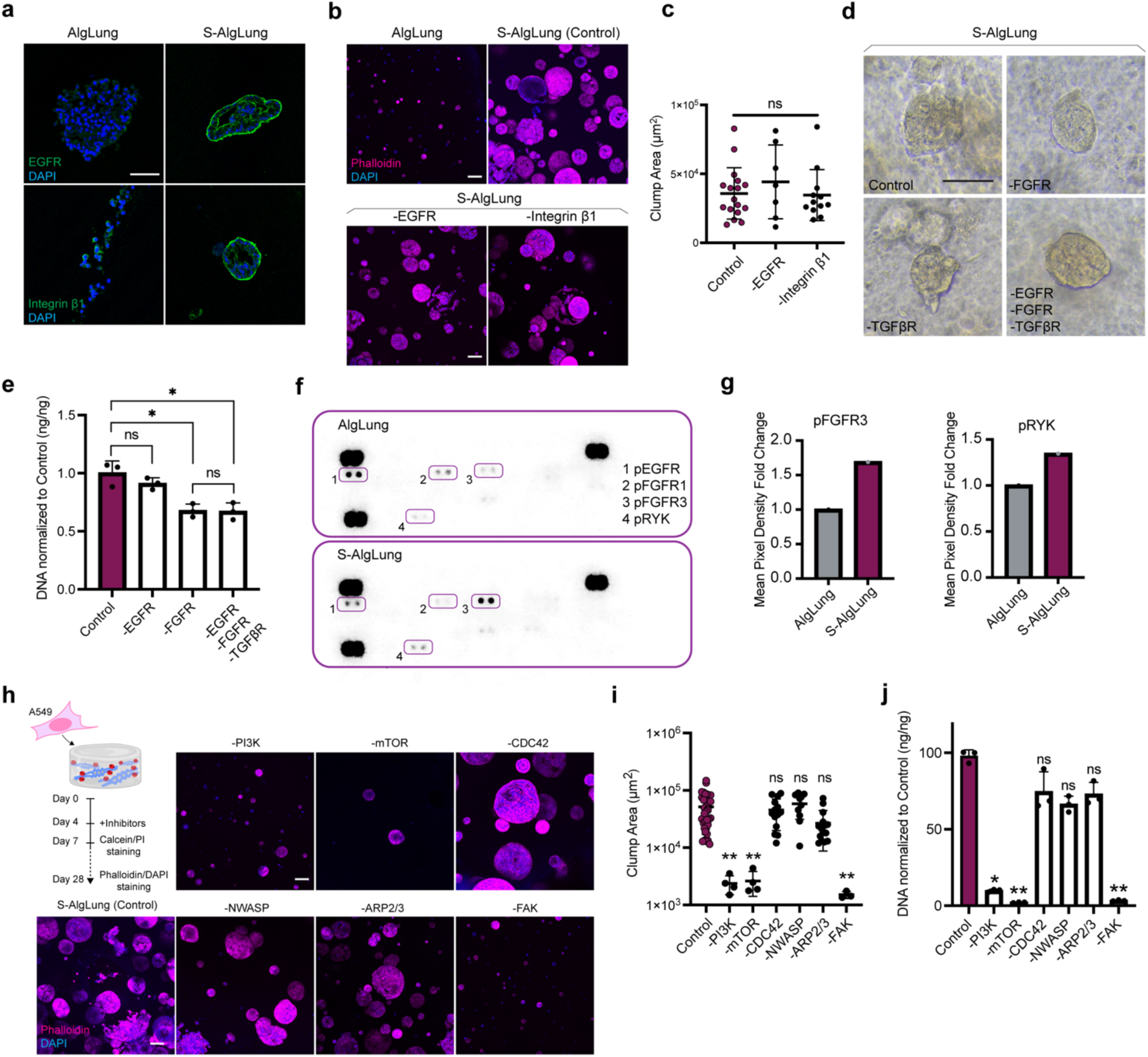
Receptor tyrosine kinase signaling mediates sulfation-induced tumorigenic phenotype. **a** Immunofluorescence staining of EGFR (top) and integrin β1 (bottom) (green) and DAPI (blue) in A549 cells grown in AlgLung and S-AlgLung hydrogels (scale bar: 100 μm) **b** Representative images of phalloidin (magenta) and DAPI (blue) stained A549 cells in AlgLung and S-AlgLung hydrogels treated with EGFR and integrin β1 inhibitors (scale bar: 100 μm). **c** Quantification of cluster area (μm^2^) of A549 cells in S-AlgLung hydrogels treated with EGFR and integrin β1 inhibitors, ns not significant **d** Representative brightfield images of A549 cells grown in S-AlgLung and treated with receptor tyrosine kinase (RTK) inhibitors at day 14, (scale bar: 100 μm). **e** Quantification of DNA content of A549 cells in S-AlgLung hydrogels and treated with RTK inhibitors. Values were normalized to control group, ns not significant, *p < 0.05. **f** Human Phospho-RTK Array performed on A549 cells grown in AlgLung and S-AlgLung hydrogels for 21 days. RTKs were framed up and footnoted. **g** Quantification of phospho-RTK expression based on 45 min exposure. Pixel density fold change was normalized to AlgLung. **h** Representative images of phalloidin (magenta) and DAPI (blue) stained A549 cells S-AlgLung hydrogels treated with indicated inhibitors (scale bar: 100 μm) **i** Quantification of cluster area (μm^2^) of A549 cells in S-AlgLung hydrogels treated with indicated inhibitors, ns not significant, **p < 0.01. **j** Quantification of DNA content of A549 cells grown in S-AlgLung hydrogels and treated with indicated inhibitors. Values were normalized to control group, ns not significant, *p < 0.05, **p < 0.01. Quantitative data is represented as mean ± S.D and statistical significance was analyzed using ordinary one-way Anova analysis.

### PI3K is a key regulator of sulfation-induced tumorigenic phenotype

Next, we focused on PI3K since it’s been reported to have a regulatory role in lung tumors and its interaction with FAK has been well-documented^44–45^. Additionally, PI3K signaling was shown to induce melanoma growth in mice upon dietary supplementation of chondroitin sulfate^17^. To validate the role of PI3K in sulfation-mediated events, we generated PIK3CA-knockdown A549 cells (Supplementary Fig. 10). PI3K loss caused a significant decrease in cell growth and cluster area in S-AlgLung hydrogels (Fig. 5a, b). Furthermore, sulfation-induced upregulation of EMT-regulating transcription factors was lost (Fig. 5c). Expression of E-cadherin was unchanged while N-cadherin was downregulated. On the other hand, fibronectin and vimentin did not show a decrease (Supplementary Fig. 11). Stemness markers SOX2, OCT3/4 and CD133 were significantly downregulated upon PIK3CA knockdown in S-AlgLung hydrogels (Fig. 5d). Thus, loss-of-function assessments revealed PI3K as an important regulator in the sulfation-induced growth, EMT and stemness on cells. To further confirm the involvement of PI3K, we performed gain-of-function experiments. We generated a PIK3CA-overexpressing A549 line (Supplementary Fig. 12) and encapsulated in healthy-mimetic AlgLung hydrogels to investigate whether PI3K overexpression alone could mimic the effect of sulfation. In AlgLung hydrogels, PIK3CA overexpressing cells proliferated rapidly during the culture period, demonstrating the importance of this pathway in lung tumor cell growth (Fig. 5e, f). Interestingly, even though proliferation was induced in AlgLung hydrogels, cellular morphology was quite different compared to sulfated hydrogels. PIK3CA-overexpressing cells displayed sheet-like growth, distinct from the cluster formation observed in S-AlgLung hydrogels (Fig. 5e). Contrarily, the expression of EMT regulators which were upregulated upon sulfation were unchanged upon PIK3CA overexpression in AlgLung hydrogels (Fig. 5g). In contrast, SLUG expression was significantly upregulated indicating that different transcription factors contribute to EMT in both contexts. Stemness markers were all significantly increased in AlgLung hydrogels demonstrating that PI3K activation alone was enough to induce stemness in lung tumor cells even in the absence of a sulfated microenvironment (Fig. 5h). Next, we focused on the role of FAK in the PI3K-mediated effects of ECM sulfation. For this, we encapsulated PIK3CA-overexpressing cells in S-AlgLung hydrogels and treated them with an FAK inhibitor. Interestingly, PIK3CA-overexpressing cells demonstrated cluster formation in sulfated hydrogels as opposed to their behavior in AlgLung, however, FAK inhibition had no effect on cell growth (Fig. 5i). Furthermore, expression of EMT markers were either not affected or increased when FAK was blocked indicating that PIK3CA overexpression compensated for FAK inhibition (Fig. 5j). These findings suggest that sulfation-promoted signaling follows the FAK-PI3K-Akt-mTOR axis in lung tumor cells (Fig. 5k).

**Figure 5.**
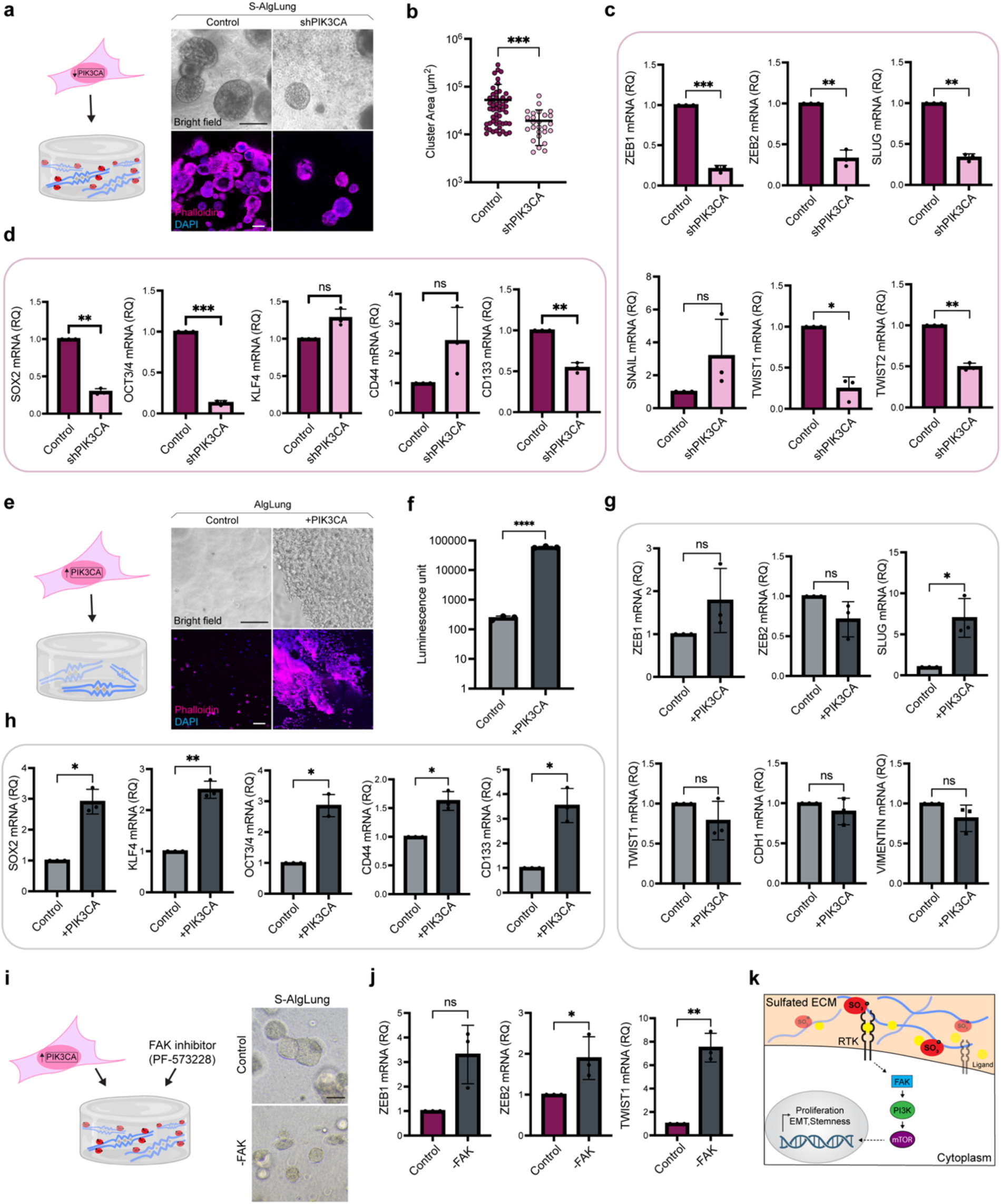
PI3K is a key regulator of sulfation-induced tumorigenic phenotype. **a** Representative brightfield and confocal microscopy images of A549 cells expressing shPIK3CA or control vectors in S-AlgLung hydrogels. Cells were stained with phalloidin (magenta) and DAPI (blue), (scale bar: 100 μm). **b** Quantification of cluster area (μm^2^) of shPIK3CA-expressing A549 cells in S-AlgLung hydrogels compared to control, ***p < 0.001. **c** mRNA expression of EMT regulators in shPIK3CA-expressing A549 cells grown in S-AlgLung hydrogels, ns not significant, *p < 0.05, **p < 0.01, ***p < 0.001. **d** mRNA expression of stemness markers in shPIK3CA-expressing A549 cells grown in S-AlgLung hydrogels, ns not significant, **p < 0.01, ***p < 0.001. **e** Representative brightfield and confocal microscopy images of A549 cells overexpressing PIK3CA or control vectors in AlgLung hydrogels. Cells were stained with phalloidin (magenta) and DAPI (blue), (scale bar: 100 μm) **f** Cell viability analysis of PIK3CA-overexpressing A549 cells in AlgLung hydrogels using CellTiter-Glo 3D assay, ****p < 0.0001. **g** mRNA expression of EMT regulators in PIK3CA-overexpressing A549 cells grown in AlgLung hydrogels, ns not significant, *p < 0.05. **h** mRNA expression of stemness markers in PIK3CA overexpressing A549 cells grown in AlgLung hydrogels, *p < 0.05, **p < 0.01. **i** Representative brightfield images of PIK3CA-overexpressing A549 cells grown in S-AlgLung hydrogels and treated with FAK inhibitor (scale bar: 100 μm). **j** mRNA expression of EMT regulators in PIK3CA-overexpressing A549 cells grown in S-AlgLung hydrogels and treated with FAK inhibitor, ns not significant, *p < 0.05, **p < 0.01. **k** Schematic illustration of the sulfated ECM-induced signaling cascade in lung tumor cells grown in S-AlgLung hydrogels. Sulfated ECM exerts affinity to bioactive ligands that leads to the activation of FGFR3 and RYK receptors and their downstream signaling. PI3K acts as a hub in sulfation-induced proliferation, EMT activation and stemness phenotype in A549 cells. All quantitative data is represented as mean ± S.D and statistical significance was analyzed using an unpaired, two-tailed student’s t-test. In qRT-PCR data, relative quantification (RQ) was used with normalization to control group.

### Sulfated ECM significantly alters the transcriptional program of cancer cells

To elucidate a bigger picture of the molecular alterations that take place in response to increased sulfation in the ECM, we profiled transcriptomes of cells grown in AlgLung and S-AlgLung hydrogels with RNAseq. Principal component analysis showed unbiased separation of clusters (Supplementary Fig. 13). We found 483 differentially expressed genes (DEGs) of which 277 genes were up-regulated and 206 were down-regulated in S-AlgLung compared to AlgLung (Fig. 6a, b). Among the DEGs, strong upregulation of SNAIL and MUC5B was observed. DEGs were enriched in a variety of biological processes involved in multiple cancer types, cell cycle, ECM-receptor interactions, cytoskeleton organization and mucin-type glycan biosynthesis (Supplementary Fig. 14). This demonstrated a collective alteration in these pathways consistently with our results on the effect of sulfation on proliferation, invasion, stemness, RTK activation and mucin expression. Understanding the causal connections between gene expression programs and cellular signaling pathways requires systems-based integrative approaches^46–47^. The precise mechanism through which alterations in ECM rewire signaling pathways remains poorly elucidated, particularly in the context of omics data analysis. Therefore, we combined transcriptional data analysis and network reconstruction to reveal the intermediate signaling mediators and signaling alterations induced by sulfated ECM. We successfully inferred the regulatory relationships between the DEGs and their corresponding transcription factors in sulfation-induced phenotype through statistical analysis and stringent filtering (see Methods) and found 34 computationally identified transcription factors (Supplementary Table 4). We then reconstructed a signaling network by orienting it from experimentally identified receptors, FGFR3 and RYK, as well as mediators, PIK3CA and PTK2 (FAK), to the inferred set of transcription factors that regulate the DEGs (Fig. 6c). Reconstructed network revealed a more complete picture of the signaling events through adding the intermediate effectors besides the hits from the RNAseq data. Network analysis identified the interaction of RYK receptor with beta-catenin (CTNNB1) which play a crucial role in lung cancer stem cell phenotype^43^, in line with our experimental data demonstrating its elevated expression in S-AlgLung hydrogels (Fig. 3f). Moreover, MYC and MYCN, located downstream of PIK3CA in the reconstructed network, are known to be deregulated in lung cancer^48^. Activated MYC is an inducer of EMT which can interact with regulators including SNAIL and TWIST. Furthermore, MYC and PI3K-Akt signaling pathway have synergistical effect in enhancing tumor growth^48^. Gene set enrichment analyses on sulfated ECM-induced network revealed pathways regulating carcinogenesis of many cancer types, cell cycle, PI3K-Akt signaling and most interestingly proteoglycan signaling (Fig. 6d). Intriguingly, when we compared the gene expression profile of PG+ patients in the TCGA LUAD cohort for the 483 DEGs between S-AlgLung and AlgLung hydrogels, PG+ patient tumors clustered with S-AlgLung, further supporting the ability of our engineered sulfated hydrogels in representing PG-rich *in vivo* tumors (Fig. 6e).

We then analyzed the expression of intermediate nodes, revealed from the regulatory network reconstruction, in PG+ patients. The heatmap indicates differential regulation of all nodes in PG+ tumors compared to normal tissues among which genes such as CDK1, CCNA2 and SRC, critical regulators of cell cycle, EMT, and progression in lung tumors^49–51^, are highly upregulated (Fig. 6f, Supplementary Fig.15). Expression of CDKN1A (Cyclin Dependent Kinase Inhibitor 1A) is repressed by MYC^52^. Our transcriptomic analysis showed that CDKN1A was differentially downregulated in cells grown in S-AlgLung hydrogels (Fig. 6b). In line with this, PG+ patient tumors showed upregulation of CDK1 (Cyclin Dependent Kinase 1) in comparison to normal tissues (Fig. 6f, Supplementary Fig. 16). PTEN, a tumor suppressor and an antagonist of PI3K signaling^53^, is another intermediate node in the network whose expression was significantly downregulated in PG+ tumors (Fig. 6c, f, Supplementary Fig. 16). Interestingly, the only intermediate which did not show any change in PG+ patient tumors compared to healthy tissues was EGFR (Supplementary Fig. 16), supporting our experimental data which revealed robustness of sulfation-mediated growth and invasiveness against EGFR inhibition (Fig. 3b-d).

**Figure 6.**
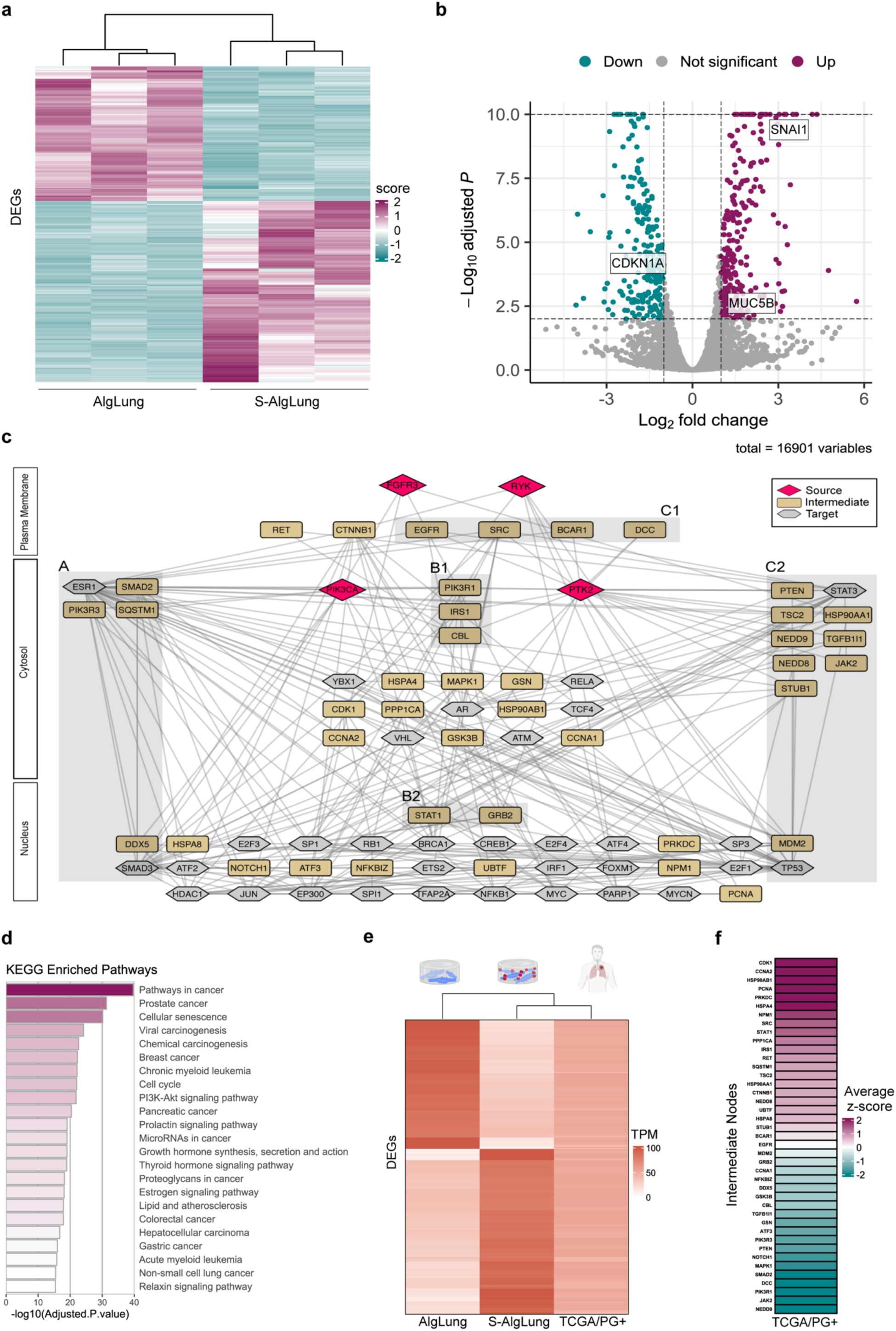
Sulfated ECM significantly alters the transcriptional program of cancer cells. **a** Heatmap shows hierarchically clustered normalized expression values of differentially expressed genes across samples. Negative z-scores are in blue color-scale, and positive z-scores are in purple color-scale. **b** Volcano plot shows DEGs in purple (up-regulated genes), blue (down-regulated genes), and gray (other genes). Thresholds to find DEGs (adj-pvalue<0.01 and abs(log2(FC))>1) are shown as black dashed horizontal and vertical lines. **c** Reconstructed signaling network orients from receptors to significant TFs that regulate the differentially expressed genes. In this network, RYK, FGFR3, PTK2 and PIK3CA are source nodes and significant TFs are target nodes. Pathlinker is used for network reconstruction. **d** Bar plot shows the functionally enriched KEGG pathways of intermediate nodes. P-values were determined using a hypergeometric test. **e** Heatmap shows TPM-normalized values of 483 DEGs, comparing TCGA/PG+ samples with S-AlgLung and AlgLung samples. Hierarchical clustering reveals that patient samples are more closely related to S-AlgLung samples.. **f** Average z-scores of mRNA expression data of intermediate nodes in TCGA/PG+ samples. Negative z-scores are in blue color-scale, and positive z-scores are in purple color-scale.

## Discussion

Lung cancer is a grave disease orchestrated by a complex series of molecular events in the TME. Emerging evidence suggest that alterations in ECM characteristics influence malignant transformation and progression^2^. There is a need for engineered models which can recapitulate the key aberrant changes in tumor matrices for elucidation of underlying signaling mechanisms^2^. Our findings reveal that expression of PGs is highly elevated among LUAD patients and correlates with invasive molecular programs including EMT and stemness. To mimic the increased PG and sGAG content in the TME, we developed a bioengineered human lung tumor model which allows tunability of sulfation within organotypic ECM. The use of decellularized native organ-derived ECMs offers the advantage of representing tissue-specific matrix onto which malignant characteristics can be introduced in a controlled manner. This is particularly important for modeling the aberrant changes in cell instructive ECM ligands considering the undefined composition of tumor-derived rBM materials. Native matrices allow representation of the ECM at the tumor’s site of origin as well as at site of metastasis using secondary organ-derived matrices. It has been established that controlling ECM ligand density is vital when modeling mechanical changes such as stiffness in engineered tumor tissues^26^. Similarly, control of mechanical properties is also very important when tuning aberrant biochemical content. Thus, our model allows recapitulation of sGAG increase in tumors while enabling independent control of ECM content, stiffness, viscoelasticity and porosity. In this study, we used a stiffness range representing healthy lung tissue to focus on the sole effect of increased sulfation. However, our model allows future investigation of synergy between sulfation and stiffening, particularly since PGs can interact with mechanosensitive receptors to activate distinct signaling mechanisms^28^. Similarly, other aberrantly increased ECM ligands in tumors such as tenascin and fibronectin families^9^ as well as cell adhesive peptide sequences^54^ can be incorporated in our model for further complexifying. Interestingly, sulfation alone induced remodeling in the ECM with elevated deposition of fibronectin and vimentin by lung tumor cells in our study. Moreover, expression of mucin-type glycans, crucial regulators of invasiveness and stemness in cancers^39^, were stimulated upon sulfation.

Our findings demonstrate that sulfation state of the cellular microenvironment is a regulator of growth and invasive phenotype marked by activation of EMT and stemness in lung cancer cells. sGAG-mimetic alginate sulfate leads to activation of a specific set of RTKs (FGFR3 and RYK) in tumor cells and the downstream FAK-PI3K-Akt-mTOR signaling axis. RTKs represent an important focus of targeted therapies, and compensatory activation of RTKs can contribute to therapeutic resistance^55^. Combination therapies that target multiple RTKs as well as ECM remodeling enzymes such as heparanase can increase the efficacy of targeted therapies^56^. Therefore, tumor-mimetic models which allow investigation of complex RTK signaling can serve as a reliable preclinical platform for developing efficient treatment strategies. Transcriptomic analyses further confirmed that distinct molecular alterations occur in response to sulfation. Our study employed network reconstruction of the transcriptomic landscape of tumor cells in engineered hydrogels. This way, transcriptomic data was integrated with experimentally identified source nodes which predicted intermediate effectors and regulatory signaling pathways that supported our hypothesis and findings. Among the signaling pathways enriched in S-AlgLung hydrogels are pathways in involved in carcinogenesis, PI3K-Akt signaling and most interestingly proteoglycan signaling which exhibits that our engineered model successfully mimicked the PG-rich microenvironment of native tumors. Additionally, clustering of PG+ LUAD patient tumors with the transcriptomic profile of S-AlgLung poses an interesting aspect. Patient tumors represent a heterogenous cast of cells, whereas our model entailed a single LUAD-derived cell line. This further emphasizes the importance of tumor ECM sulfation in determining the transcriptional programs which modulate growth and invasiveness. The fact that our enrichment analyses hit not only NSCLC but also many different types of cancers (prostate, gastric, breast, myeloid leukemia) underlines that sulfation is a critical pan-cancer ECM characteristic and suggests that our model can be expanded to different tumors. Lastly, these approaches which entail tunability and customization of engineered tumor models can be adapted to patient-derived organoids for testing therapeutics as a further step towards precision medicine.

## Materials & Methods

### Bioinformatics Analyses for Publicly Available Patient Data

The cBioPortal database was used to download the human lung cancer data of Lung Adenocarcinoma (TCGA, PanCancer Atlas), and Lung Adenocarcinoma (CPTAC, Cell 2020 datasets)^57–58^. From the Lung Adenocarcinoma (LUAD) TCGA datasets, mRNA expression z-scores relative to normal samples (log RNA Seq V2 RSEM) dataset was utilized (Fig. 1a). A list of proteoglycan genes was obtained from the Matrisome Database^59^, with 34 genes being present in the TCGA dataset (Supplementary Table 2). The pairwise correlations among these genes were visualized in a heatmap using ComplexHeatmap (v2.13.1) in R (https://cran.r-project.org/package=BiocManager, https://www.R-project.org). Highly expressed proteoglycan gene names were identified, and correlation values were hierarchically clustered both by rows and columns. Samples were divided into two groups. A sample was considered proteoglycan positive (PG+) if the z-score expression value was equal to or higher than the cut-off of three for at least three genes, while a sample was considered proteoglycan negative (PG-) if the z-score expression value was equal to or higher than the cut-off but the number of genes is equal to zero. Proteoglycan, EMT and CSC scoring was done considering the proteoglycan gene list and literature-curated EMT and CSC gene lists, respectively (Supplementary Table 2). Scores were derived via computation of the average expression of genes in the relevant list for each patient sample. A comparison between the proteoglycan scores of PG+ and PG-samples was conducted to assess significant differences between the two groups (Fig. 1c). The result was shown as a box plot generated with the help of ggpubr of R (v0.6.0) (https://cran.r-project.org/package=ggpubr, https://www.R-project.org). Two regression plots were generated to visualize the relationship between EMT scores and Proteoglycan scores with the same samples on both TCGA LUAD transcriptomic and CPTAC LUAD proteomic data using the proteoglycan and EMT gene list in Supplementary Table 2 (Fig. 1e,f). For the CPTAC LUAD data, the average of the z-scores of protein abundance ratios was used to calculate the EMT and PG scores with the same gene lists mentioned. Regression plots were created using Python’s seaborn (v0.12.0) (https://pypi.org/project/seaborn/). Spearman rank correlation was employed to calculate correlation values. TCGA LUAD clinical patient and clinical sample datasets were used to conduct a survival analysis (Fig. 1d). The ggfortify package in R (v0.4.14) is used to generate the survival plot (https://cran.r-project.org/web/packages=ggfortify, https://www.R-project.org).

### sGAG Quantification

5 NSCLC LUAD samples and their normal parenchyma counterparts (Supplementary Table 3) were collected with Koc University Institutional Review Board (2020.001.IRB2.001) ethics approval and consent of participants undergoing lobectomy as part of their clinical care. Quantification of sGAG content was performed using Blyscan Sulfated Glycosaminoglycan Assay kit (Biocolor, UK) following manufacturer’s instructions. Briefly, weighted tumor and normal parenchyma samples were digested in 125 µg/ml papain (Sigma) buffer (400 mg sodium acetate, 200 mg EDTA and 40 mg cysteine in 50 ml of 0.2 M sodium phosphate buffer, pH 6.4) at 65°C overnight. Samples were then mixed with dye reagent followed by dye retrieval and absorbance measurement at 656 nm using a microplate reader. sGAG content in tumor and normal parenchyma samples were normalized to their wet weight.

### Tissue Histology

NSCLC LUAD tumor and matched normal parenchyma tissue samples were fixed with 3.7% formaldehyde solution (EMS) at 4°C overnight, followed by immersion in 30% sucrose overnight. Samples were then embedded in OCT (Tissue-Tek), snap-frozen, sectioned as 10 µm slices and mounted on glass slides. For haematoxylin & eosin staining, slides were hydrated and stained with Mayer’s Haematoxylin (Merck) for 3 min followed by a 3-min wash with tap water. Then, slides were immersed in 95% ethanol and stained with Eosin solution (Bright-Slide) for 45 seconds. To visualize deposition of sGAGs in samples, slides were hydrated and stained with 1% Alcian Blue (Sigma) in 3% acetic acid solution (pH 2.5) for 30 min followed by a 2-min wash with tap water. After staining, all slides were dehydrated with graded alcohol treatments, mounted, and visualized by light microscopy.

### Modification of Alginate

Sulfation of alginate (Novamatrix) was carried out as previously described^22^. Briefly, chlorosulfonic acid (HClSO_3_) (Sigma) was diluted in formamide (Sigma) and added dropwise onto alginate while stirring. The reaction was carried out at 60°C with agitation for 2.5 h. Alginate sulfate was precipitated with cold acetone and re-dissolved in ultra-pure water and neutralized overnight. The solution was purified by dialyzing in 12 kDa molecular weight cut-off (MWCO) dialysis tubing (Sigma) against 100 mM NaCl for 48 hours and ultra-pure water for 72 hours with solution change every 12 hours and then lyophilized.

### Chemical Characterization of Alginate Sulfate

Elemental analysis of sulfur content in alginate sulfate was performed using high-resolution inductively coupled mass spectrometry (ICP-MS). Degree of sulfation (DS), number of sulfate groups per monomer, was estimated from the mass balance equation assuming one sodium counterion for each negatively charged group and one water molecule per monosaccharide: Monosaccharide mass = C_6_O_6_H_5_ + (DS+1) Na^+^ + (DS) SO_3_^−^ + H_2_O. Molecular weight of alginate sulfate was analyzed by size exclusion chromatography with a multiangle laser light detection system (SEC-MALS) using a refractive index (dn/dc) of 0.15 for all samples.

### Decellularization and characterization of lung tissues

Decellularization of bovine lung was carried out as previously described^21^. Briefly, lung tissue pieces were thoroughly washed in dH_2_O with 2% Penicillin/Streptomycin/Amphotericin (P/S/A) and then subjected to freeze-thaw cycles using liquid nitrogen. Next, tissue pieces were treated with 10U/ml DNase (Sigma), washed again in dH_2_O, lyophilized and, cryomilled into a powder form. dECM powder was then digested in 1 mg/ml pepsin solution (Sigma) at a final concentration of 15 mg/ml (w/v) at room temperature for 48 hours, neutralized, lyophilized, and stored at -20°C for further use. To validate elimination of cellular content in decellularized bovine lung (dLung), samples were fixed in 3.7% formaldehyde solution, embedded in OCT, cryo-sectioned and mounted on glass slides. Haematoxylin & Eosin and Hoechst staining was performed as previously described^21^. To characterize collagen and sGAG content, Sirius red and Alcian blue stainings were performed, respectively, as previously described^21^.

### Cell Culture

Human lung adenocarcinoma cell line A549 (#CCL-185) was purchased from American Tissue Culture Collection (ATCC) and cultured in growth medium (DMEM/F12 (Lonza) supplemented with 10% FBS (Biowest) and 1% PenStrep (Gibco)). Human embryonic kidney (HEK293T) cells were cultured with DMEM High Glucose (Biowest) supplemented with 10% FBS and 1% P/S. Both cell lines were maintained in an incubator at 37 °C and 5% CO_2_ and tested for Mycoplasma using MycoAlert Mycoplasma Detection Kit (Lonza) regularly.

### Hydrogel Formation and Cell Encapsulation

Alginate and alginate sulfate were dissolved in serum-free DMEM/F12 and sterile filtered using 0.2 µm pore size syringe filter. Lyophilized dLung was also dissolved in serum-free DMEM/F12 containing 2% P/S overnight at 4°C. Calcium sulfate stock solution was prepared in ddH_2_O and autoclaved. Before the cell encapsulation in hydrogels, A549 cells were expanded as monolayer cultures, trypsinized, centrifuged, counted and resuspended in growth medium. The final cell density in hydrogels was 5*10^4^ ml^-1^ unless stated otherwise. Constituents were combined using a double-syringe and Luer lock coupler system and hydrogels were deposited into a 24-well plate. Hydrogels were allowed to solidify for 45 min in an incubator before adding culture medium. Alginate-Matrigel (Growth Factor Reduced, Corning) (AlgMat) hydrogels were prepared following a similar approach.

### Mechanical Characterization

Mechanical characterization of hydrogels was performed using a Discovery HR-2 rheometer (TA instruments). Briefly, hydrogels were deposited onto the lower plate which was pre-cooled to 4°C and a 20 mm parallel plate was lowered until the gap reached 1 mm. Mineral oil (Sigma) was applied at the periphery of the gels to prevent dehydration during the measurement. Oscillatory rheology was used to measure the storage modulus at constant frequency and amplitude (1 Hz, 1% strain) for 2 hours. For viscoelasticity measurements, creep-recovery test was performed with application of a constant shear stress for 1 h on gels while strain was recorded. Then, the sample was unloaded, and strain was measured for 2 hours. All measurements were done at least in triplicates.

### Hydrogel Porosity Assessment

Rhodamine-tagged dextran (Invitrogen) was encapsulated into AlgLung and S-AlgLung hydrogels at a concentration of 0.5 mg/ml. Two different molecular weights(10 kDa/70 kDa) were used for dextran. Hydrogels were cast on a 24-well plate and incubated in PBS for 3 days. Media was removed every 24 hours for measuring the diffusion of rhodamine. Quantification of released dextran was performed with fluorescence readout using a microplate reader at 570 nm/590 nm excitation/emission. At least three replicates were used for each condition.

### Quantification of DNA

dsDNA quantification from hydrogels was performed by Quant-iT PicoGreen dsDNA Assay Kit (Invitrogen) according to manufacturer’s instructions. Briefly, at designated time points, hydrogels were collected, washed once with wash buffer (150 mM NaCl and 5 mM CaCl_2_) and stored at −80 °C until the assay was carried out. Hydrogels were digested in 125 µg/ml papain buffer containing 10 mM EDTA, 100 mM sodium phosphate, 100 mM sodium acetate, 10 mM L-cysteine (pH 6.4) at 60°C overnight. After digestion, diluted samples were mixed with Quant-iT PicoGreen reagent and incubated for 5 min at room temperature. Fluorescence was measured at 520 nm with excitation at 485 nm in a microplate reader. Experiments were done using at least three hydrogels for each group.

### CellTiter-Glo® 3D Cell Viability Assay

CellTiter-Glo® 3D Cell Viability Assay (Promega) was performed in hydrogels according to manufacturer’s instructions with minor modifications. Hydrogels were briefly washed and incubated with assay buffer for 45 min at room temperature after a 5-min shake in a plate shaker. Luminescence was measured for at least 3 hydrogels using a microplate reader.

### Assessment of Cell Viability and Morphology

A549 cells encapsulated in hydrogels were stained at the beginning (Day 7) and the end of culture (Day 28) with 2 µM Calcein-AM (Invitrogen) and 30 µg/mL propidium iodide (PI) (Sigma) in growth medium to assess viability. To monitor the morphological changes of encapsulated A549 cells in hydrogels, samples were fixed with 4% PFA for 30 min at room temperature. The gels were then washed with wash buffer three times for 5 min followed by permeabilization and blocking in 5% BSA (Sigma), 1% Triton X-100 (Sigma) for 1 hour at room temperature. F-actin staining was performed using Phalloidin-iFluor 488 Reagent (1:1000, Abcam) for 45 min at room temperature followed by a 5-min wash. Nuclei staining was performed with DAPI (1 µg/mL, Sigma) and samples were washed twice with wash buffer for 10 min before imaging. Samples were imaged with Leica SP8 confocal microscope and z-stacks were obtained with a 5 μm step length for at least 3 hydrogels.

### Cluster Analysis

Cluster area and invasiveness of clusters formed in hydrogels which were stained for F-actin and nuclei were analyzed using ImageJ. For each condition, at least 3 hydrogels were imaged and at least 5 z-stack images were analyzed. Briefly, z-stack images were projected with maximum intensity, thresholded, then boundaries of clusters were calculated using the ‘’analyze particles’’ module. In some images, clusters were overlapped making it hard to analyze the individual clusters. In these cases, z-stack slices were individually analyzed, and total cluster numbers were normalized to total stack number. Invasiveness of clusters were calculated using the circularity output from the cluster size analysis. A circularity value of 1 obtained from analysis indicates a perfect circular cell cluster which shows 0% invasiveness whereas a value of 0 circularity indicates 100% invasiveness.

### Immunofluorescence

Hydrogels were fixed with 4% PFA for 30 min at room temperature. Next, the gels were washed with wash buffer three times for 5 min followed by permeabilization and blocking in 5% goat serum (Gibco), 1% BSA, 1% Triton X-100 for 1 h at room temperature. Primary antibody incubation was performed overnight at 4°C in staining buffer containing 1% BSA, 0.1% Triton X-100. Primary antibodies used for these studies are listed in Supplementary Table 5. Hydrogels stained with primary antibodies were washed twice for 10 min at room temperature with staining buffer. Secondary antibody incubation was performed overnight at 4°C using goat anti-rabbit FITC (1:200, H&L, Jackson ImmunoResearch) or goat anti-mouse Alexa Fluor 633 (1:200, H&L, Invitrogen) followed by 2 hours wash with staining buffer at room temperature. Nuclei staining was performed with 1 µg/mL DAPI in staining solution followed by a 15-min wash with staining buffer. Samples were imaged with Leica SP8 confocal microscope.

### Phospho-RTK Array

Phosphorylation of RTKs in A549 cells encapsulated in hydrogels were revealed by Proteome Profiler Human Phospho-RTK Array (R&D) according to manufacturer’s instructions. Briefly, AlgLung and S-AlgLung hydrogels were treated with cell retrieval buffer (100 mM Sodium Citrate, 50 mM EDTA in ddH_2_O, pH: 7.2) to retrieve cells from the hydrogels. After two washing steps for 5 min with ice-cold PBS, cell lysates were collected by lysis buffer provided in the kit and quantified with BCA Protein Assay. Arrays were incubated with 100 µg of total protein for both conditions. Images were captured by LI-COR imaging system Odyssey Fc with Image Studio Acquisition Software. The quantification of each dot was performed using the longest exposed images by ImageJ and the expression of p-RTKs in S-AlgLung hydrogels were normalized to AlgLung samples.

### Inhibition Assays

Drug inhibitors with the indicated concentrations in Supplementary Table 6 were added in culture medium of hydrogels at day 4 and renewed every 3 days until day 21. Same amount of DMSO was also added in culture medium as a control group. At the end of the experiment, 3 hydrogels were kept for DNA quantification in each group, and the remaining hydrogels were fixed and stained for F-Actin as previously described in the text to analyze cluster morphologies.

### Plasmid Constructs

Full length plasmid of PI3KCA (phosphatidylinositol-4,5-biphosphate 3-kinase catalytic subunit alpha, NM_006218.4) was purchased from Addgene (Plasmid ID: 81736, Hahn and Root Lab). The coding sequence of PI3KCA was cloned into Lenti-PCDH-EF1-mNeonGreen-MCS-T2A-puromycin plasmid (a kind gift from Fırat-Karalar Lab, Koç University, Istanbul). GFP-tagged shRNA plasmids targeting PI3KCA(NM_0062) and scrambled shRNA plasmid were obtained from Vector Builder (plasmid IDs: VB9000558918GSY, VB9000619992CNJ and VB9000558929 respectively).

### Restriction Enzyme Cloning and Primers

PI3KCA open reading frame was cloned into PCDH-mNeonGreen backbone with restriction-dependent cloning. Phusion High Fidelity DNA Polymerase (NEB) was used with the following primers: PI3KCA_human_forward_BamH1 5′-ACTGGGATCCATGCCTCCACGACCATCATC-3′; PI3KCA_human_reverse_Not1: 5′-ACTGGCGGCCGCGTTCAATGCATGCTGTTTAATTGTG-3′.

The PCR reaction was performed and the resulting PCR product as well as PCDH-mNeonGreen-MCS-T2A-puromycin plasmid were cleaved with BamH1(NEB) and Not1(NEB) enzymes. Digested products were then ligated with T4 DNA Ligase (NEB).

### Lentiviral Production and Stable Line Generation

HEK293T cells were transfected with Lipofectamine 3000 Transfection Reagent (Invitrogen) with a ratio of 4:3:1 for Lentiviral DNA, psPAX2, PCMV-VSVG respectively. 48 hours post-transfection viral supernatants were collected and concentrated with PEG8000 (Sigma). A549 cells were transduced with lentiviral particles expressing the gene of interest with exogenous fluorescent tag with a MOI of 5. For enhancing the infection efficiency, 10 µg/mL of Protamine Sulfate (Sigma) was used. Cells were selected with 1.5 µg/mL Puromycin (Sigma) for 5 days.

### Sorting of PI3KCA-overexpressing Cells

A549 cells transduced with mNeonGreen-PI3KCA viral plasmids were sorted utilizing Attune NxT Flow Cytometer to establish a homogenous PI3KCA-overexpressing population. Cells were trypsinized and centrifuged at 300g for 5 minutes. Supernatant was removed carefully and pellet was resuspended with Magnesium and Calcium free PBS supplemented with 3% BSA. Cell suspension was run through a strainer with 40 µm mesh size to obtain a single cell suspension. FSCA vs SSC and FSCH vs FSCA gates were applied to separate single cells and eliminate doublets. Then, mNeonGreen-expressing transduced cells were sorted with FITCA vs FSCH.

### Western Blotting

Cells were collected in PBS (Biowest) and centrifuged at 300 g for 3 min. Supernatant was discarded. Cell pellet was lysed with RIPA buffer (EcoTech) supplemented with PhosSTOP (Roche) and cOmplete Mini-EDTA Free (Roche) for 40 min on ice. The protein concentration was determined with BCA Protein Assay (Pierce). 50 µg protein was loaded in each well of precast 4-15% Bis Tris gels (BioRad) and run for 2 hours at 80V. Wet transfer was performed onto a PVDF (BioRad) membrane for 2 hours at 110V. Membrane was washed with Tris-Buffered Saline with Tween-20 (TBST) for 3 times. Blocking was performed with 5% non-fat dry milk (BioRad) and incubated at 4°C overnight. Membrane was washed with TBST for 3 times. Primary antibodies were prepared as follows: PI3KCA (p110a) (#A0265, Abclonal) at 1:500, GAPDH (#AB9485, Abcam) at 1:1000 in TBST with 5% BSA (Sigma) and 0.02% Sodium Azide (Sigma) and incubated at 4°C overnight. Membrane was washed with TBST for 3 times. Secondary antibody was prepared as follows: Rabbit IgG H&L (HRP) (#ab97051, Abcam) at 1:10000 dilution in TBST with 5% non-fat milk and incubated for 45 min at room temperature. After incubation, membranes were washed with TBST and incubated with a highly sensitive ECL solution (Pierce) for chemiluminescence detection. Visualization was performed with LI-COR imaging system Odyssey Fc with Image Studio Acquisition Software.

### Real Time-Quantitative Polymerase Chain Reaction (qRT-PCR)

RNA extraction from hydrogels was performed with TRIzol Reagent (Invitrogen). After adding TRIzol, the gels were crushed with a pestle on dry ice, then centrifuged at 12000 rpm for 5 min at 4°C. Chloroform was added to the supernatant and incubated on ice for 10 min followed by centrifugation at 12000 rpm for 15 min at 4°C. Aqueous phase was collected and equal volume 70% EtOH was added. From this point, RNA isolation was performed with Nucleospin RNA II (MN) kit following manufacturer’s instructions. 1 µg RNA was reverse-transcribed using M-MLV Reverse Transcriptase Kit (Invitrogen). qRT-PCR was performed using SYBR Green (Roche) on a LightCycler (Roche) equipment. Primer sequences that were used are given in Supplementary Table 7.

### Transcriptomic Analyses and Network-based Data Integration

RNA sequencing was performed using DNBSEQ Platform by BGI Genomics (Hong Kong). The quality of the reads was checked using FastQC and mapped against the ENSEMBL Homo Sapiens reference genome (GRCh38) using STAR version 2.7.3a^60^. Counts were calculated from the aligned reads using the featureCounts function of the Rsubread R package^61^. Differential analysis and normalization were performed using the DESeq2 R package and R (version 4.2.2)^62^. Genes with more than 10 reads were used in the DESeq2 analysis. Additional filtering of genes with extreme count outliers and low mean normalized counts was performed by DESeq2, yielding 16901 genes. For the representation of gene abundance transcripts per million (TPM) values were computed using DGEobj.utils R package. Genes were considered differentially expressed when log2 fold changes were equal or greater than 1.0 for upregulated genes or equal or lower than -1.0 for downregulated genes, with a Benjamini–Hochberg p-adjusted value less than or equal to 0.01. For functional enrichment analysis of GO Biological Process and KEGG Datasets, EnrichR^63^ was used. For gene-set enrichment analysis for GO Biological Process, WebGestalt^64^ was used. A signaling network was reconstructed using PathLinker version 1.4.3^65^ in Cytoscape version 3.8.2^66^ where iRefWeb was selected as the reference interactome. Protein-protein interactions that have a confidence score of less than 0.4 were filtered out from the iRefWeb interactome. Additionally, UBC and its interactions were deleted. Filtered iRefWeb interactome was used as the reference interactome, “undirected” and “unweighted” parameters were selected, and iterations were made with the given source and target nodes with a k value of 1000. Obtained subnetwork (226 nodes, 754 edges) was used as a reference, “undirected”, “weights are probabilities=confidence”, and “edge penalty=1” parameters were selected and iterations were made without changing source and target nodes with a k value of 50000. After this iteration, a subnetwork (77 nodes, 258 edges) was obtained. The network was visualized with information on the source, target, and subcellular locations.

## ACKNOWLEDGEMENTS

Authors acknowledge funding from the International Fellowship for Outstanding Researchers Program of TÜBİTAK (118C238), European Union’s Horizon 2020 research and innovation program under the Marie Skłodowska-Curie grant agreement 101032602, and Swiss National Science Foundation (P2EZP2**-**172172). The entire responsibility of the publication belongs to the owner, the financial support from TÜBİTAK does not mean that the content of the publication is approved in a scientific sense by TÜBİTAK. BioRender.com was used to create the icons in Figures 2a, 3c, and 5. The authors gratefully acknowledge Dr. Stephen Spinella for helping with the preliminary mechanical characterization experiments. The authors gratefully acknowledge the use of services and facilities of Koç University Research Center for Translational Medicine (KUTTAM).

