## Supplementary Information for "Extracellular matrix sulfation in the tumor microenvironment stimulates cancer stemness and invasiveness"

#### Supplementary Figures

##### Supplementary Figure S1

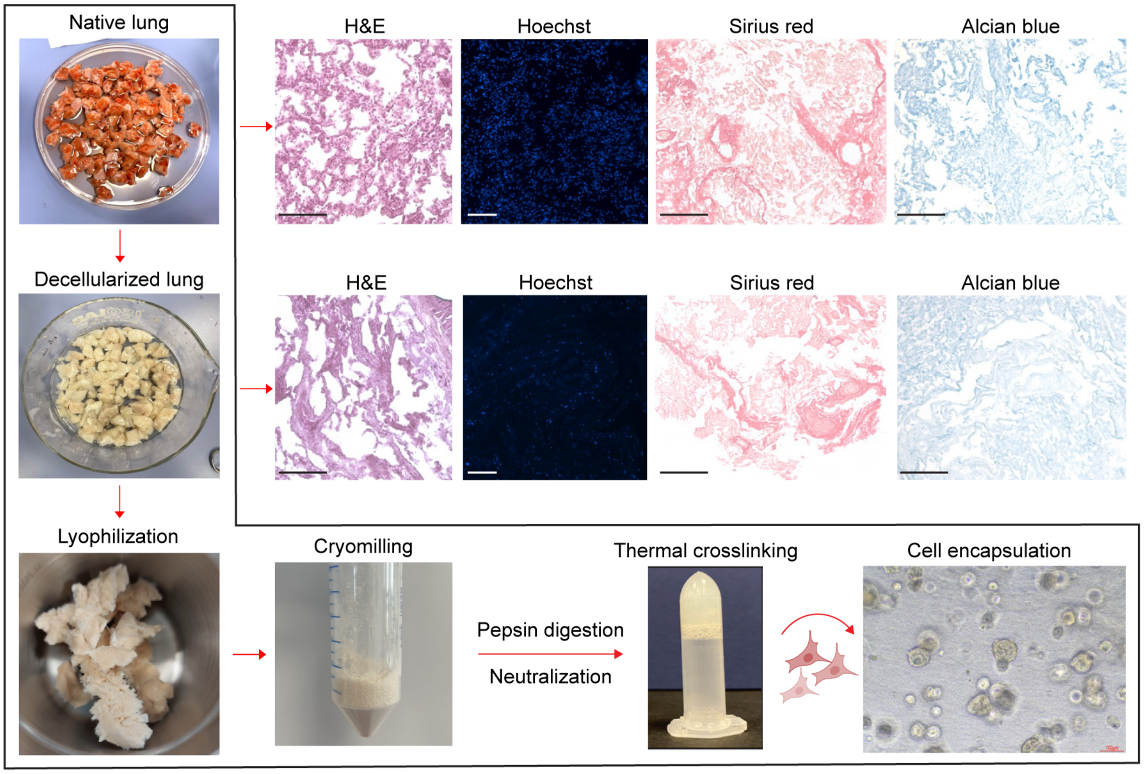


**Supplementary Figure S1. Steps of bovine lung decellularization and characterization of decellularized tissue.** Native bovine lung was cut into small pieces, washed throughly with $ddH_2O$ supplemented with 1% Penicillin/Streptomycin/Amphotericin (P/S/A) followed by freeze-thaw cycles in liquid nitrogen. Decellularized lung tissue pieces were lyophilized, cryomilled into a powder form and digested in pepsin solution (1 mg/ml in 0.01 M HCl, pH:2) for 48 hours at room temperature. After pepsin digestion, pre-gel solutions were neutralized using NaOH and buffered to neutral pH. A549 cells were encapsulated into dLung hydrogels and monitored for cellular growth via brightfield microscopy. Elimination of nuclear content in decellularized lung tissues was shown by Haematoxylin & Eosin as well as Hoechst stainings in comparison to native tissue. Collagen and sGAG retention in decellularized lung tissues were revealed by Sirius Red and Alcian Blue stainings, respectively (Scale bar: 100 µm).

**Supplementary Figure S2**

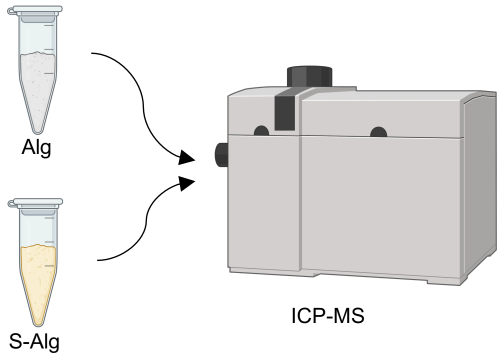


| Sample | S | | Degree of Sulfation (DS) |
|---|---|---|---|
| | mg/kg | RSD (%) | Sulfate groups/monomer |
| Alg | 621 | 3,2 | 0 |
| S-Alg | 46 698 | 2,3 | 0.41 |

**Supplementary Figure S2. Analysis of sulfate content in S-Alg compared to Alg.** Elemental analyses were performed with high-resolution inductively coupled plasma mass spectrometry (ICP-MS) for determining sulfur content in Alg and S-Alg samples (RSD: Relative Standard Deviation).

**Supplementary Figure S3**

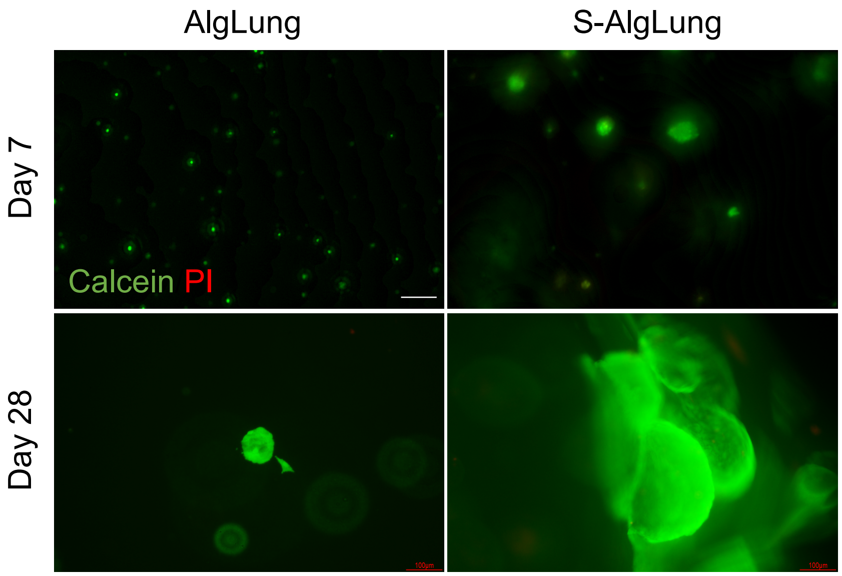


**Supplementary Figure S3. Viability assessment of A549 cells grown in AlgLung and S-AlgLung hydrogels.** Calcein AM (green) and propidium iodide (PI) (red) staining of A549 cells grown in AlgLung and S-AlgLung hydrogels at day 7 and day 28 (Scale bar: 100 μm).

**Supplementary Figure S4**

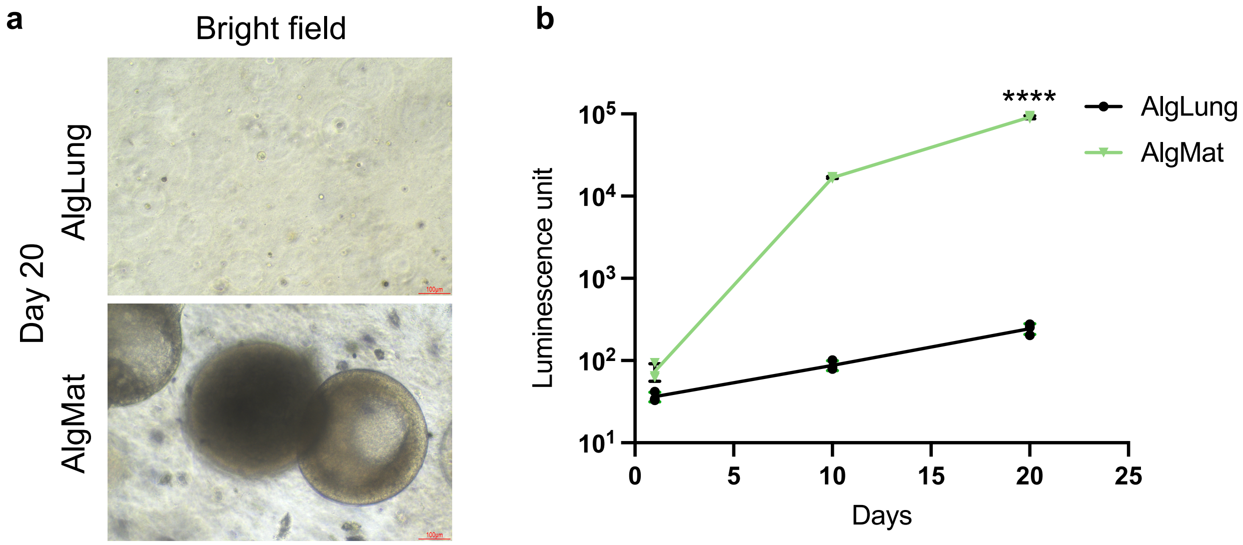


**Supplementary Figure S4. Growth of A549 cells in AlgMat hydrogels. a** Representative brightfield images of A549 cells grown in AlgLung and AlgMat (Alginate-Matrigel) hydrogels at day 20 (Scale bar: 100 µm). **b** Cellular growth analysis by CellTiter-Glo 3D assay in AlgLung and AlgMat hydrogels. Data is represented as mean ± S.D and statistical significance is analyzed using an unpaired, two-tailed student's t-test, ****p < 0.0001.

**Supplementary Figure S5**

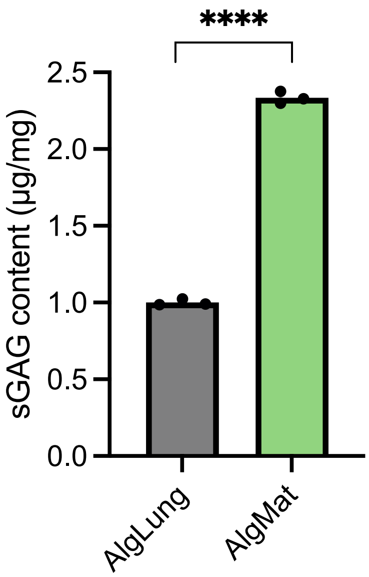


**Supplementary Figure S5. Assessment of sGAG content in AlgLung and AlgMat hydrogels**. Data is represented as mean ± S.D and statistical significance is analyzed using an unpaired, two-tailed student's t-test, **** indicates $p<0.0001$.

#### Supplementary Figure S6

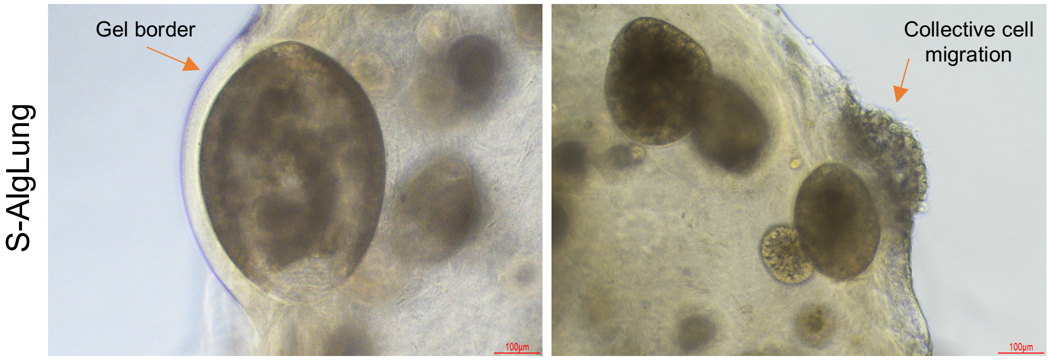


**Supplementary Figure S6. Sulfated hydrogels induce collective cell migration.** A549 clusters grown in S-AlgLung hydrogels displayed collective cell migration towards the periphery of the hydrogels, shown via brightfield microscopy images (Scale bar: 100 μm).

**Supplementary Figure S7**

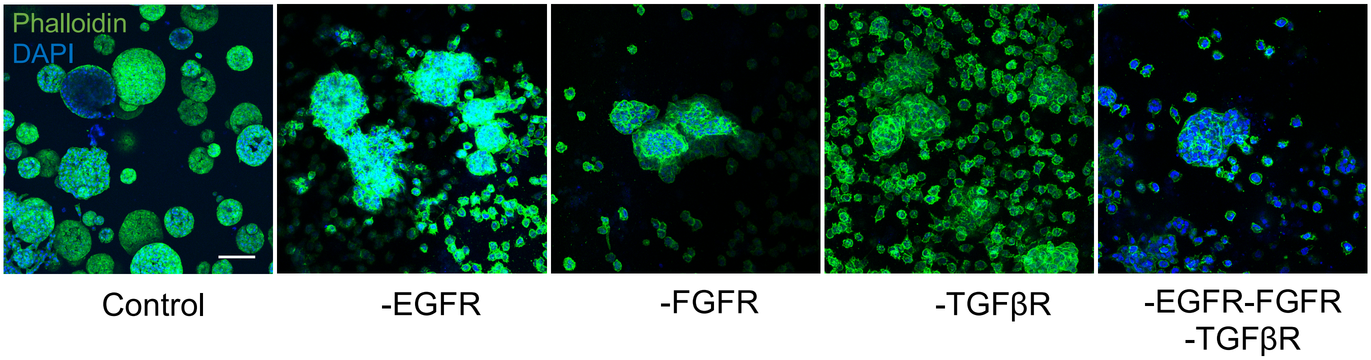


**Supplementary Figure S7.** Representative immunofluorescence images of Phalloidin (green) and DAPI (blue) stained A549 cells in S-AlgLung hydrogels treated with inhibitors targeting indicated proteins (Scale bar: 100 μm).

**Supplementary Figure S8**

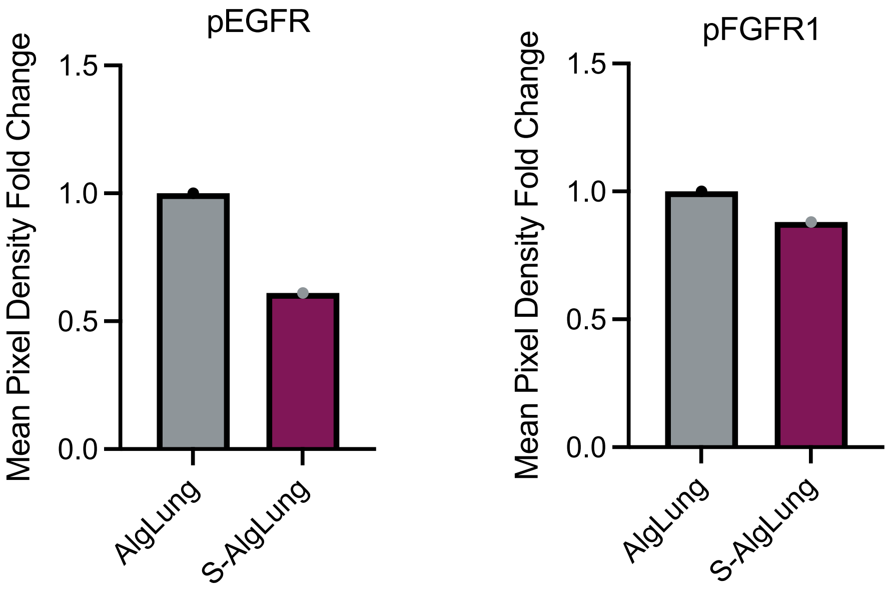


**Supplementary Figure S8. Phosphorylation of EGFR and FGFR1.** The quantitation of phosphorylated EGFR and FGFR1 expression from phospho-proteome array (45 min exposure). Pixel density fold change was represented normalized to AlgLung.

#### Supplementary Figure S9

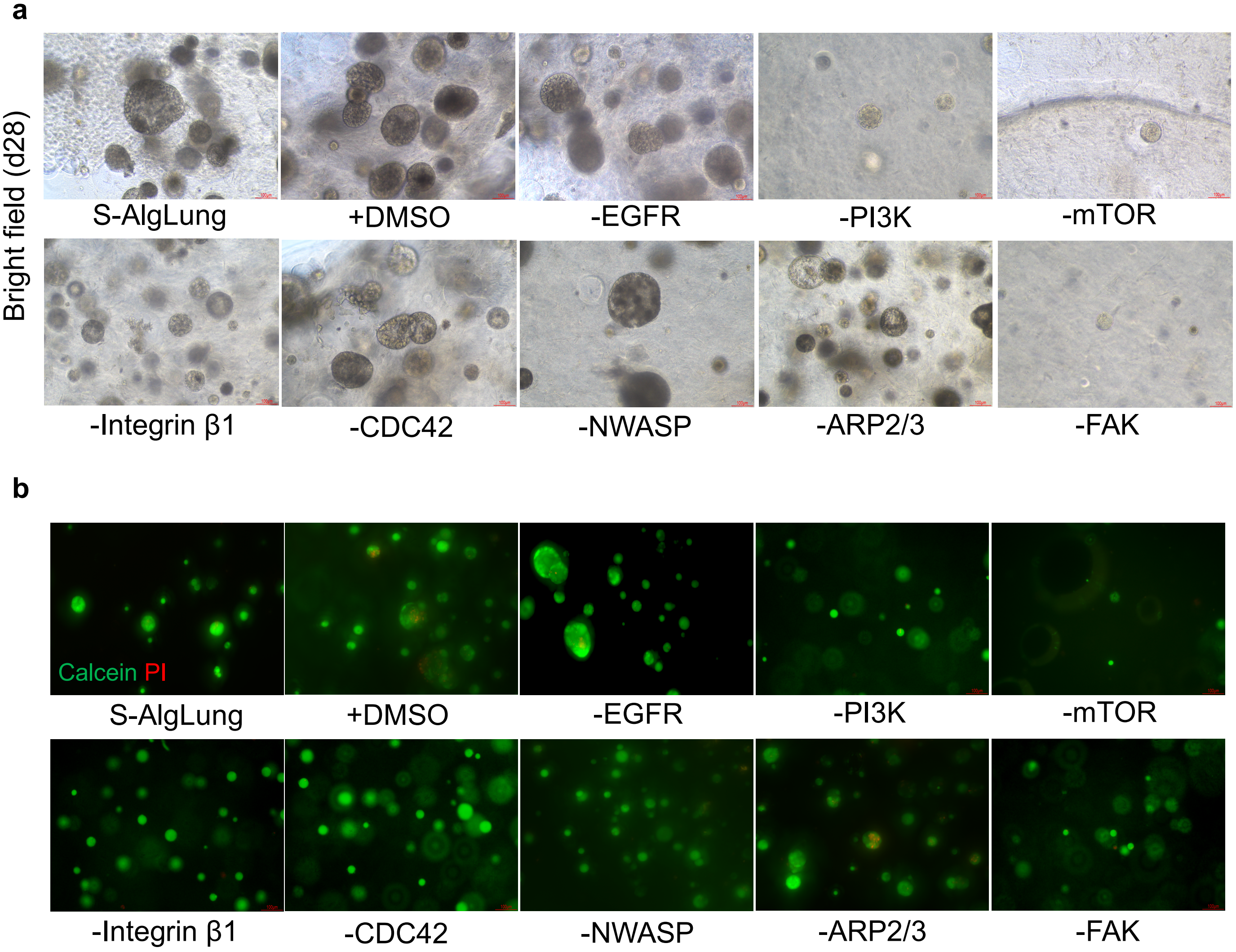


**Supplementary Figure S9. Inhibitor screens.** Representative **a** brightfield and **b** immunofluorescence images of calcein AM (green), propidium iodide (red) stained A549 cells in S-AlgLung hydrogels treated with DMSO or inhibitors targeting indicated proteins (Scale bar: 100 μm).

**Supplementary Figure S10**

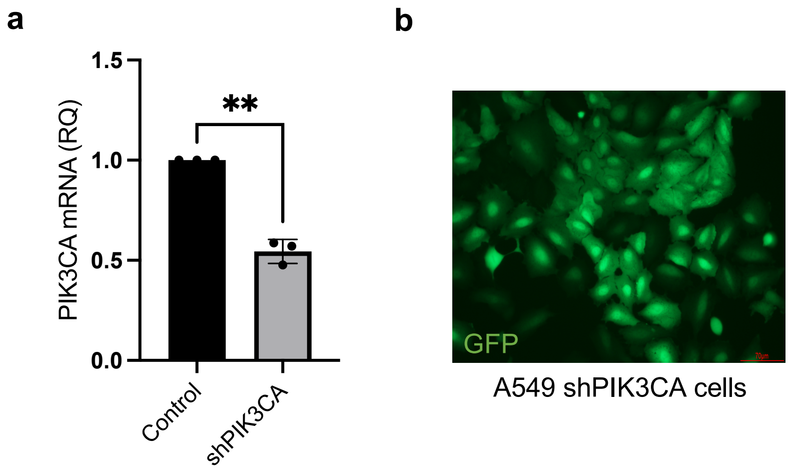


**Supplementary Figure S10. Validation of PI3K knockdown. a** mRNA expression of PIK3CA in shPIK3CA expressing A549 cells. Data is represented as mean ± S.D and statistical significance is analyzed using an unpaired, two-tailed student's t-test, ***p < 0.001. **b** Representative images of shPIK3CA expressing A549 cells tagged with GFP (Scale bar: 70 μm).

**Supplementary Figure S11**

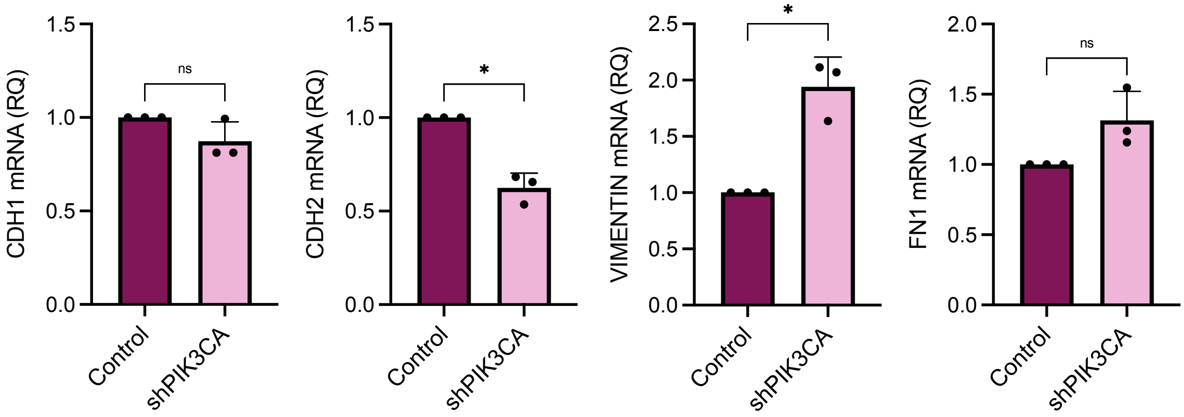


**Supplementary Figure S11. Gene expression of EMT markers in knockdown A549 cells.** mRNA expression of E-cadherin, N-cadherin, vimentin and fibronectin in shPIK3CA-expressing A549 cells grown in S-AlgLung hydrogels. Relative quantification (RQ) was used with normalization to control group. Data is represented as mean ± S.D and statistical significance is analyzed using an unpaired, two-tailed student's t-test, ns not significant, *p < 0.05.

#### Supplementary Figure S12

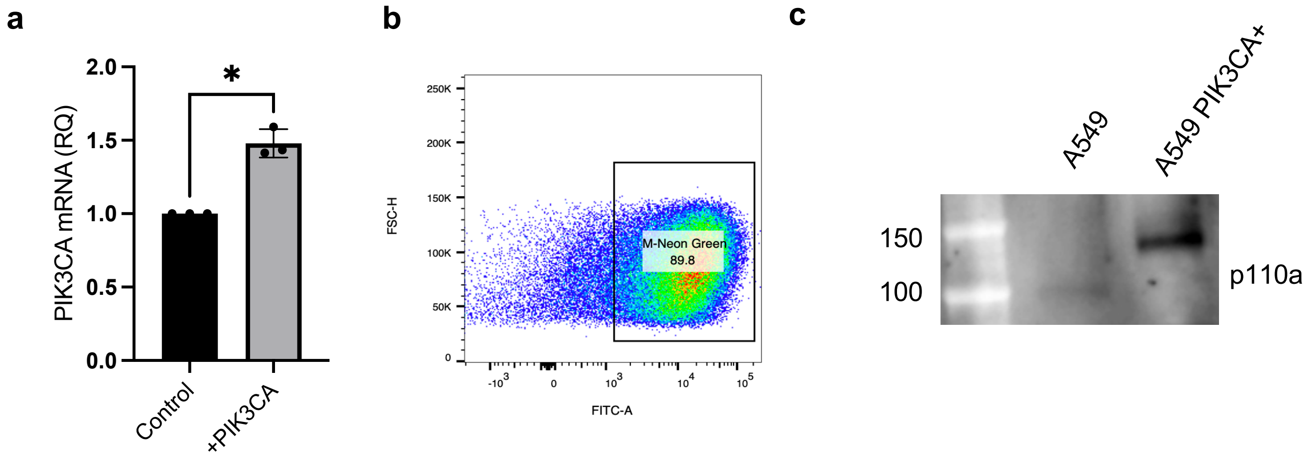


**Supplementary Figure S12. Validation of PIK3CA-overexpressing A549 cell line. a** PIK3CA mRNA expression of PIK3CA-overexpressing A549 cells. Data is represented as mean ± S.D and statistical significance is analyzed using an unpaired, two-tailed student's t-test, *p<0.05. **b** FACS plot demonstrating sorting for M-Neon Green-tagged PIK3CA-overexpressing A549 cells. **c** Validation of PIK3CA overexpression through assessment of p110a protein expression in A549 cells by western blotting.

#### Supplementary Figure S13

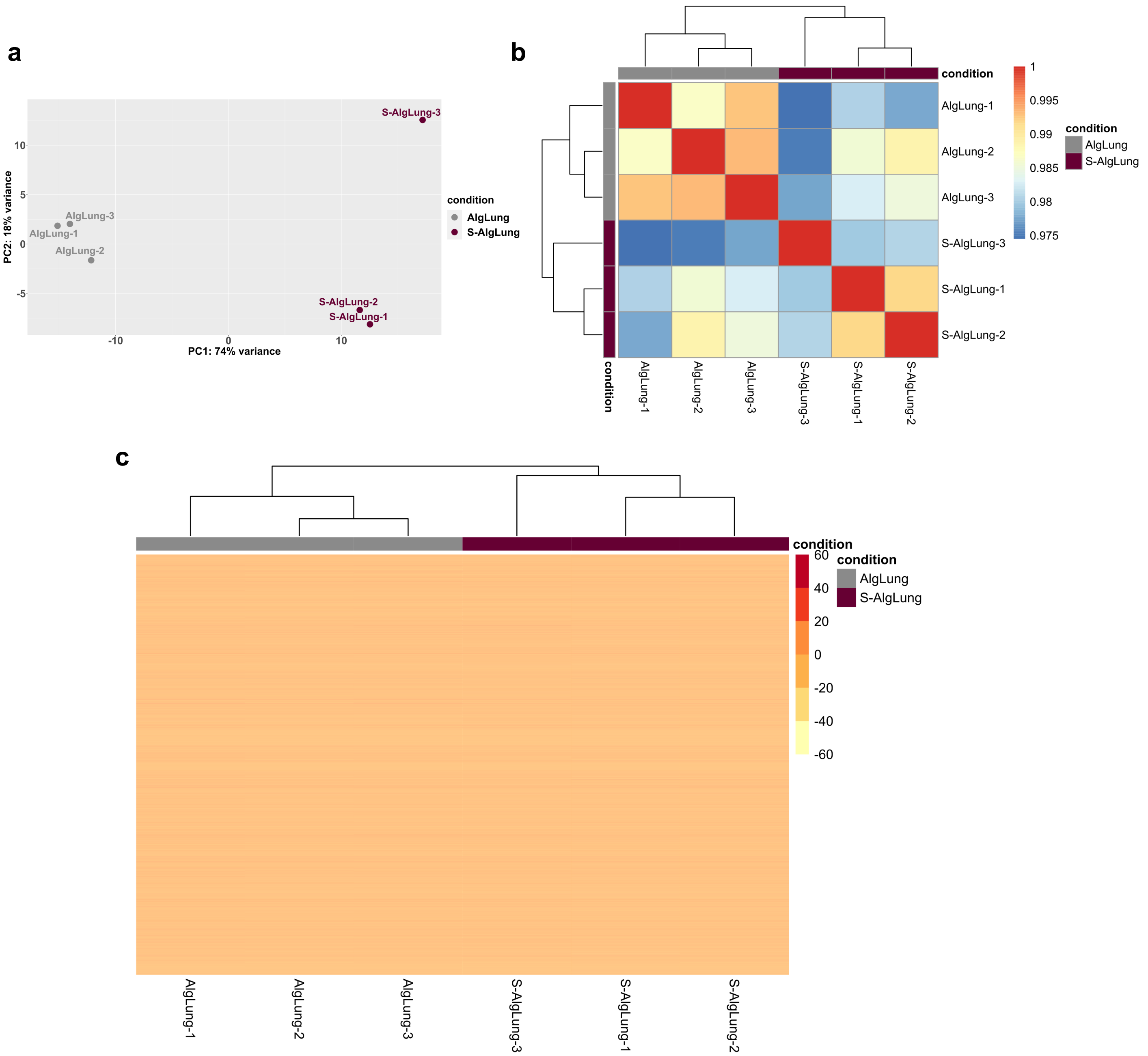


**Supplementary Figure S13. a** Principal component analysis (PCA) showed a clear separation of AlgLung from S-AlgLung hydrogels. **b** The correlation heatmap shows the Pearson correlation between samples, where samples of the same condition exhibit higher correlation values. **c** Heatmap shows DESeq2's median of ratios normalized counts for all genes (16,901).

**Supplementary Figure S14**

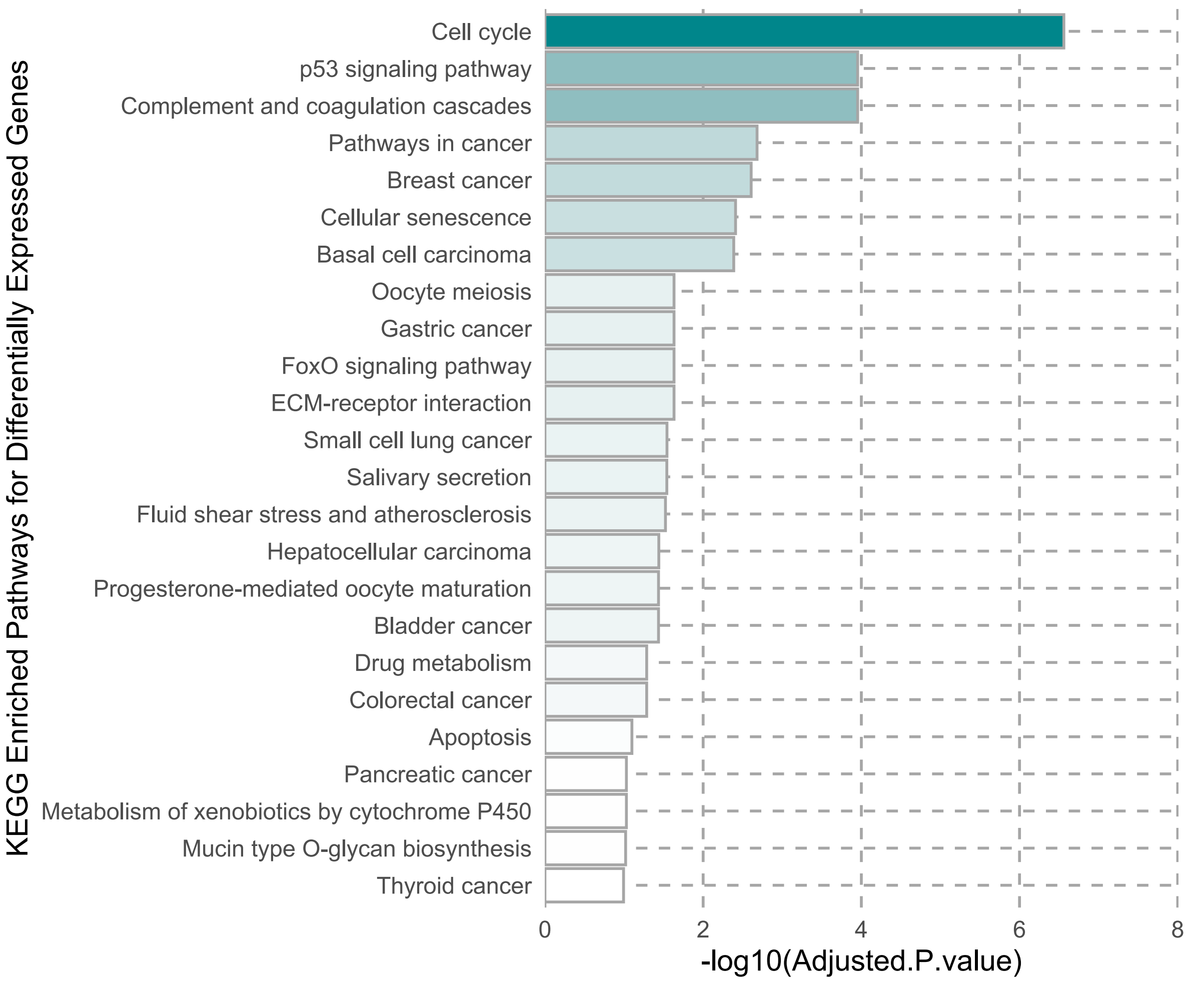


**Supplementary Figure S14.** Bar plots showing the functional enrichment of KEGG pathways in up-regulated and down-regulated genes in S-AlgLung hydrogels.

#### Supplementary Figure S15

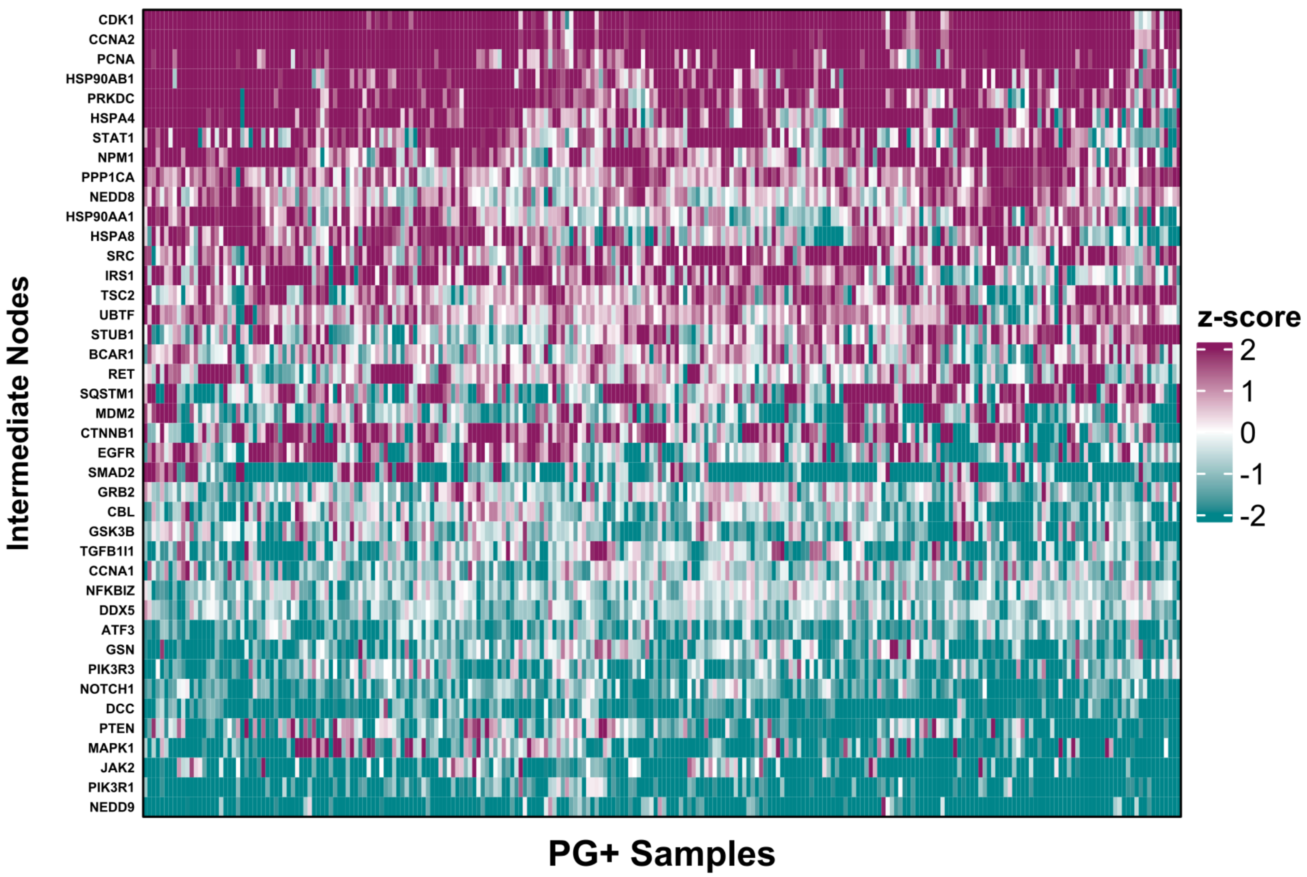


**Supplementary Figure S15.** Heatmap shows intermediate genes in PG+ TCGA LUAD patient tumor samples. Each column represents a patient.

#### Supplementary Figure S16

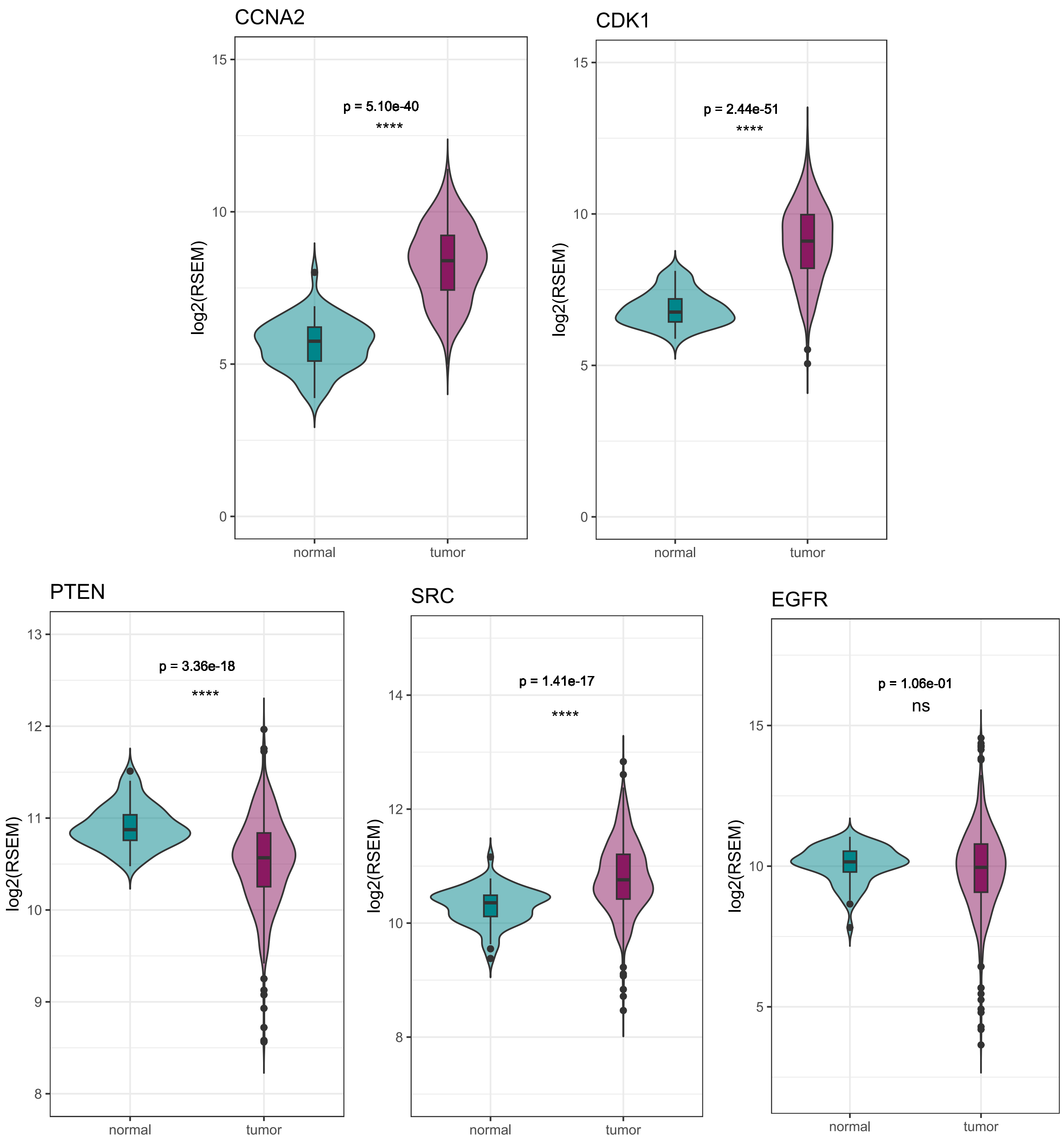


**Supplementary Figure S16.** Expression profiles of CCNA2, CDK1, PTEN, SRC and EGFR in TCGA LUAD patient tumor samples compared to normal lung tissue. Statistical analyses were performed using a two-sample t-test. ns indicates non-significant, **** indicates p<0.0001.

#### Supplementary Tables

##### Supplementary Table 1

| | Alg | | | S-Alg | | |
|---|---|---|---|---|---|---|
| $M_n$ (kDa) | 111,7 | 107,2 | 109,5 | 70,6 | 67,1 | 68,9 |
| $M_w$ (kDa) | 215,3 | 211,7 | 213,5 | 124,2 | 124,5 | 124,4 |
| Polydispersity ($M_n$/$M_w$) | 1,9 | 2 | 2 | 1,8 | 1,9 | 1,9 |

**Supplementary Table 1.** Molecular weight of Alg and S-Alg samples revealed by size exclusion chromatography (SEC-MALS).

**Supplementary Table 2**

| Proteoglycan Genes | EMT-Related Genes | CSC-Related Genes |
|---|---|---|
| ACAN | CDH1 | CD24 |
| ASPN | CDH2 | CD34 |
| BCAN | FN1 | CD38 |
| BGN | VIM | CD44 |
| CHAD | DSP | CD90 |
| CHADL | OCLN | CD133 |
| DCN | MMP1 | KLF4 |
| EPYC | MMP2 | LIN28A |
| ESM1 | MMP3 | LIN28B |
| FMOD | MMP9 | MYC |
| HAPLN1 | MMP12 | NANOG |
| HAPLN2 | SNAI1 | POU5F1 |
| HAPLN3 | SNAI2 | SOX2 |
| HAPLN4 | SNAI3 | FOXP1 |
| HSPG2 | TWIST1 | NOTCH1 |
| IMPG1 | TWIST2 | NOTCH2 |
| IMPG2 | ZEB1 | DLL1 |
| KERA | ZEB2 | DLL4 |
| LUM | SPARC | DDR1 |
| NCAN | TIMP1 | DDR2 |
| NYX | SERPINE1 | DKK1 |
| OGN | | FZD7 |
| OMD | | WNT1 |
| PODN | | SMO |
| PODNL1 | | ABCB1 |
| PRELP | | ABCB5 |
| PRG2 | | ALDH1A1 |
| PRG3 | | ALDH1A3 |
| PRG4 | | ALDH3A1 |
| SPOCK1 | | BMI1 |
| SPOCK2 | | ENG |
| SPOCK3 | | KIT |
| SRGN | | AXL |
| VCAN | | FOXA2 |

**Supplementary Table 2.** Proteoglycan, EMT and and CSC-related genes used in bioinformatic analyses.

**Supplementary Table 3**

| | Age | Gender | Diagnosis |
|---|---|---|---|
| Patient #1 | 63 | Male | Lung Adenocarcinoma |
| Patient #2 | 63 | Female | Lung Adenocarcinoma |
| Patient #3 | 52 | Female | Lung Adenocarcinoma |
| Patient #4 | 69 | Female | Lung Adenocarcinoma |
| Patient #5 | 59 | Female | Lung Adenocarcinoma |

**Supplementary Table 3.** Age and gender information of patient donors whose tumor and normal lung parenchyma tissues were used to perform histological stainings and sGAG assay in Figure 1.

### Supplementary Table 4

| Key TF | Description | # of overlapped genes | P value | Q value | List of overlapped genes |
|---|---|---|---|---|---|
| TP53 | tumor protein p53 | 22 | 2.12e-11 | 1.66e-9 | BIRC5,FAS,CCNA2,CCNB2,CD82,MKI67,PLK1,ATF3,KLF2,PTTG1,GADD45A,PRC1,VEGFA,CDKN1A,RECQL4,FOXM1,EGR1,E2F1,CCNB1,DUSP1,MMP2,CDK1 |
| E2F1 | E2F transcription factor 1 | 20 | 2.42e-11 | 1.66e-9 | MYBL2,CCNB1,CDK1,E2F1,FOXM1,RRM2,CDC6,CDKN1A,AURKB,BIRC5,AURKA,PDK4,DUSP1,PLK1,KIF2C,VEGFA,DAPK2,TOP2A,ASF1B,MYCN |
| E2F3 | E2F transcription factor 3 | 8 | 7.61e-11 | 3.48e-9 | CCNB1,PLK1,CDC6,CDKN1A,AURKA,CDK1,CCNA2,VEGFA |
| NFKB1 | nuclear factor of kappa light polypeptide gene enhancer in B-cells 1 | 26 | 6.7e-9 | 2.29e-7 | PTGDS,EGR1,PDK4,CXCR4,SERPINA3,TNC,SNAI1,PSMB9,TFF3,ATF3,SOCS3,CDKN1A,CCNB1,MMP2,VEGFA,CD40,E2F1,PTAFR,MUC5B,GADD45G,B2M,POU2F2,BIRC5,CCL2,FAS,HES1 |
| YBX1 | Y box binding protein 1 | 9 | 1.36e-8 | 3.73e-7 | TOP2A,CCNB1,MMP2,E2F1,CDC20,FAS,MYCN,CXCR4,ERBB3 |
| ATF2 | activating transcription factor 2 | 9 | 2.57e-8 | 5.86e-7 | PCK1,CDKN1A,ATF3,JUN,PLAT,FAS,ITGB8,MMP2,DUSP1 |
| TFAP2A | transcription factor AP-2 alpha (activating enhancer binding protein 2 alpha) | 12 | 6.01e-8 | 1.18e-6 | CDKN1A,VEGFA,MMP2,FAS,CRABP2,L1CAM,CD82,CCNB1,ACHE,ADM,MCAM,IGFBP5 |
| SP1 | Sp1 transcription factor | 31 | 1.23e-7 | 2.11e-6 | A,UGT2B15,MYCN,GIPR,CCL2,MMP2,MYBL2,TNC,EGR1,FOXM1,FBN1,MACROD1,CA9,RECQL4,VEGFA,PTTG1,C4B,KIF2C,BIRC5,E2F1,CDKN1A,FAS,MMP11,PLAT,PADI1,CCNA2,TK1,PADI3,F7,FOS,PC |
| JUN | jun proto-oncogene | 16 | 2.83e-7 | 4.31e-6 | PLAT,CDKN1A,FAS,CCL2,MMP2,ITGB8,CD82,ATF3,ETS2,STMN1,TNC,NEFL,UGT2B15,JUN,VEGFA,LBP |
| ATM | ataxia telangiectasia mutated | 7 | 3.72e-7 | 5.1e-6 | FOXM1,FAS,PBK,CDKN1A,DUSP1,GADD45A,VEGFA |
| RELA | v-rel reticuloendotheliosis viral oncogene homolog A (avian) | 23 | 4.11e-7 | 5.12e-6 | TFF3,EGR1,VEGFA,FAS,E2F1,CD40,MMP2,PDK4,BIRC5,PSMB9,HES1,CCL2,POU2F2,TNC,CDKN1A,CXCR4,MUC5B,GADD45G,PTAFR,SNAI1,SERPINA3,CCNB1,SOCS3 |
| E2F4 | E2F transcription factor 4, p107/p130-binding | 7 | 5.25e-7 | 5.99e-6 | PCLAF,PLK1,BIRC5,TTK,CCNB1,MCM10,AURKB |
| KLF4 | Kruppel-like factor 4 (gut) | 7 | 2.86e-5 | 2.83e-4 | BIRC5,VEGFA,MMP2,ATF3,LAMA3,CCNB1,CDKN1A |
| FOXM1 | forkhead box M1 | 5 | 2.89e-5 | 2.83e-4 | CCNB1,FGB,CDC6,VEGFA,KRT15 |
| HDAC1 | histone deacetylase 1 | 9 | 3.12e-5 | 2.85e-4 | E2F1,RECQL4,EGR1,CD82,TXNIP,SFRP1,BIRC5,FOS,CDKN1A |
| BRCA1 | breast cancer 1, early onset | 8 | 4.08e-5 | 3.49e-4 | CDKN1A,FST,GADD45A,EGR1,ASPM,CCNB1,VEGFA,FOS |
| SP3 | Sp3 transcription factor | 11 | 5.2e-5 | 4.19e-4 | CA9,GIPR,MYCN,CDKN1A,SOCS3,PADI3,PADI1,PLAT,MMP2,VEGFA,BIRC5 |
| RB1 | retinoblastoma 1 | 6 | 5.97e-5 | 4.47e-4 | CDKN1A,VEGFA,FOS,E2F1,BIRC5,CDK1 |
| MYCN | -myc myelocytomatosis viral related oncogene, neuroblastoma derived (aviar | 7 | 6.32e-5 | 4.47e-4 | MCM10,CDKN1A,MYBL2,MCM2,BIRC5,MCM4,HLA-B |
| KLF6 | Kruppel-like factor 6 | 5 | 6.85e-5 | 4.47e-4 | ATF3,CDKN1A,PTTG1,DAPK2,TXNIP |
| VHL | von Hippel-Lindau tumor suppressor, E3 ubiquitin protein ligase | 5 | 6.85e-5 | 4.47e-4 | COL4A2,CA9,CXCR4,KLF10,VEGFA |
| MYC | v-myc myelocytomatosis viral oncogene homolog (avian) | 10 | 9.05e-5 | 5.64e-4 | VEGFA,CCNB1,HLA-B,ATF3,CXCR4,CCNA2,SFRP1,FOXM1,UBE2C,CDKN1A |
| STAT3 | ignal transducer and activator of transcription 3 (acute-phase response facto | 12 | 9.61e-5 | 5.73e-4 | FOS,BIRC5,CFB,MUC5B,MMP2,FGG,VEGFA,FAS,CDKN1A,SOCS3,S1PR1,CCL2 |
| PARP1 | poly (ADP-ribose) polymerase 1 | 5 | 2.14e-4 | 0.00112 | FBN1,SNAI1,CDKN1A,JUN,E2F1 |
| TCF4 | transcription factor 4 | 5 | 2.14e-4 | 0.00112 | JUN,BIRC5,WNT4,FST,VEGFA |
| AR | androgen receptor | 9 | 2.6e-4 | 0.00123 | ERBB3,CDC6,HMMR,CDKN1A,JUN,PIP,UGT2B15,KISS1R,VEGFA |
| EP300 | E1A binding protein p300 | 7 | 2.61e-4 | 0.00123 | VEGFA,CA9,BIRC5,NR0B2,CCNB2,LAMA3,CDKN1A |
| SMAD3 | SMAD family member 3 | 5 | 6.12e-4 | 0.0025 | VEGFA,TNC,JUN,FST,CDKN1A |
| ETS2 | v-ets erythroblastosis virus E26 oncogene homolog 2 (avian) | 5 | 7.12e-4 | 0.00279 | EGR1,CDKN1A,CDK1,MMP2,TNC |
| IRF1 | interferon regulatory factor 1 | 6 | 9.97e-4 | 0.00334 | CDKN1A,CDK1,PSMB9,E2F1,CD40,CCNB1 |
| CREB1 | cAMP responsive element binding protein 1 | 8 | 1 | 0.00334 | PLAT,FOS,MMP2,CXCR4,STMN1,MSMB,JUN,SNAI1 |
| ATF4 | activating transcription factor 4 (tax-responsive enhancer element B67) | 5 | 0.00108 | 0.00354 | F7,ATF3,CA9,VEGFA,CCL2 |
| ESR1 | estrogen receptor 1 | 7 | 0.00167 | 0.00519 | VEGFA,CDKN1A,E2F1,CCNA2,UGT2B15,JUN,FOS |
| SPI1 | spleen focus forming virus (SFFV) proviral integration oncogene spi1 | 6 | 0.00276 | 0.00766 | CD40,DAPK2,PRTN3,CCL2,ELANE,MACROD1 |

**Supplementary Table 4.** List of transcription factors with a Q value smaller than or equal to 0.01 and the number of overlapped genes higher than or equal to 5 retrieved from the TRRUST database using the differentially expressed genes and provided as target nodes to Pathlinker for network construction.

**Supplementary Table 5**

| Protein | Antibody |
|---|---|
| EGFR | Biolegend (352901) |
| Integrin β1/ITGB1 | SantaCruz (sc-13590L) |
| E-Cadherin | Cell Signaling Technologies (3195) |
| Vimentin | Cell Signaling Technologies (5741) |
| N-Cadherin | Cell Signaling Technologies (13116) |
| Fibronectin | Abcam (ab2413) |
| β-Catenin | Abcam (ab6302) |
| SOX2 | Biolegend (630801) |
| MUC5B | Sigma (HPA008246) |
| P63 | Cell Signaling Technologies (13109) |
| TTF-1 | Cell Signaling Technologies (12373) |

**Supplementary Table 5.** List of antibodies used in immunofluorescence.

**Supplementary Table 6**

| Target | Inhibitor | Concentration |
|---|---|---|
| EGFR | Erlotinib | 10 μM |
| FGFR | PD173074 | 500 nM |
| TGFBR | A83-01 | 500 nM |
| Integrin β1 (P5D2) | SC13590 | 1 μg/ml |
| PI3K | LY294002 | 20 μM |
| CDC42 | ML141 | 10 μM |
| ARP2/3 | CK-666 | 100 μM |
| NWASP | Wiskostatin | 10 μM |
| FAK | PF-573228 | 5 μM |
| mTOR | Rapamycin | 250 nM |

**Supplementary Table 6.** List of inhibitors and their final concentrations used in inhibition assays.

#### Supplementary Table 7

| | | | |
|---|---|---|---|
| PI3KCA_human_forward | CCATATCTCCAAAGTAGAAC | VIMENTIN_human_forward | ACCAGCTAACCAACGACAAAG |
| PI3KCA_human_reverse | TAGAATGACGACTAGCAGTA | VIMENTIN_human_reverse | AAAGATTGCAGGGTGTTTTCG |
| ZEB1_human_forward | ACCCTTGAAAGTGATCCAGC | FN1_human_forward | CAGAGGCATAAGGTTCGGG |
| ZEB1_human_reverse | CATTCCATTTTCTGTCTTCCGC | FN1_human_reverse | TTCAGACATTCGTTCCCACTC |
| ZEB2_human_forward | ACTCCTGTCTGTCTCGCAAA | SOX2_human_forward | CTTTTATGAGAGAGATCCTG |
| ZEB2_human_reverse | GCTCGATAAGGTGGTGCTTG | SOX2_human_reverse | ACCGTACCACTAGAACTTT |
| SNAIL_human_forward | GGAAGCCTAACTACAGCGAG | OCT3/4_human_forward | CACTAAGGAAGGAATTGG |
| SNAIL_human_reverse | CAGAGTCCCAGATGAGCATTG | OCT3/4_human_reverse | GTGTGTCTATCTACTGTGTCC |
| SLUG_human_forward | AGCATTTCAACGCCTCCA | KLF4_human_forward | TCTGTGACTGGATCTTCTAT |
| SLUG_human_reverse | GGATCTCTGGTTGTGGTATGAC | KLF4_human_reverse | CTTCCTCTTCTTCTAACATC |
| TWIST1_human_forward | GCATTCTCAAGAGGTCGTGC | CD44_human_forward | CTCACTCAAGCTCTTTAACT |
| TWIST1_human_reverse | TGACTATGGTTTTGCAGGCC | CD44_human_reverse | GAATATCTAGAAGGAGTGGA |
| TWIST2_human_forward | AGAGCGACGAGATGGACAAT | CD133_human_forward | ACCTACAGCATATTCTTCAC |
| TWIST2_human_reverse | ACAGACTCGAATGCATCCCA | CD133_human_reverse | CTGTTAAACTGTACACCGTA |
| CDH1_human_forward | CTGCCAATCCCGATGAAATTG | MUC5AC_human_forward | ACCAGCATCTTCATCAACCT |
| CDH1_human_reverse | TCCTTCATAGTCAAACACGAGC | MUC5AC_human_reverse | AAGTCATTAACGGCGATGTC |
| CDH2_human_forward | CAGAATCAGTGGCGGAGATC | MUC5B_human_forward | CTACATCAAGGTCAGCATCCG |
| CDH2_human_reverse | CAGCAACAGTAAGGACAAACATC | MUC5B_human_reverse | ATAGAACTCGTTGAAGGCCG |
| GAPDH_human_forward | CTGACTTCAACAGCGACACC | GAPDH_human_reverse | GTGGTCCAGGGGTCTTACTC |

**Supplementary Table 7.** List of primers used in RT-qPCR.
